# The synaptonemal complex aligns meiotic chromosomes by wetting

**DOI:** 10.1101/2024.08.07.607092

**Authors:** Spencer G. Gordon, Alyssa A. Rodriguez, Yajie Gu, Kevin D. Corbett, Chiu Fan Lee, Ofer Rog

## Abstract

During meiosis, the parental chromosomes are drawn together to enable exchange of genetic information. Chromosomes are aligned through the assembly of a conserved interface, the synaptonemal complex, composed of a central region that forms between two parallel chromosomal backbones called axes. Here we identify the axis-central region interface in *C. elegans*, containing a conserved positive patch on the axis component HIM-3 and the C-terminus of the central region protein SYP-5. Crucially, the canonical ultrastructure of the synaptonemal complex is altered upon weakening this interface. We developed a thermodynamic model that recapitulates our experimental observations, indicating that the liquid-like central region can assemble by wetting the axes without active energy consumption. More broadly, our data show that condensation drives tightly regulated nuclear reorganization during sexual reproduction.

## Introduction

Cellular processes are tightly controlled spatially, requiring that large structures, such as organelles or chromosomes, be moved and precisely positioned. This is most commonly achieved by motor proteins and the polymerization/depolymerization of cytoskeletal filaments. These active processes consume free energy provided by the hydrolysis of nucleotide triphosphate (NTP) molecules to move cargo over large distances. However, an alternative mechanism that could regulate cellular organization has been proposed: thermodynamically-driven formation of protein assemblies (Brangwynne *et al*. 2009). Biomolecular condensates interact with membrane-bound organelles and other large cellular structures (Gouveia *et al*. 2022), and synthetic condensates are capable of exerting pico-newton-scale forces on adjacent structures (Strom *et al*. 2024). However, the functional importance of condensate assembly for moving cellular structures *in vivo* has not been demonstrated.

A cellular structure whose maneuvering is particularly well-regulated is the chromosome. During meiosis, the specialized cell division cycle that produces gametes, the unassociated homologous parental chromosomes (homologs) are brought together and aligned along their lengths (Zickler and Kleckner 2023). Paired and aligned homologs are necessary for the formation of exchanges (crossovers) that shuffle the maternal and paternal genomes and allow chromosomes to correctly segregate into the gametes. Errors in these intricately controlled processes lead to aneuploidy, congenital birth defects and infertility.

Chromosome alignment in meiosis is driven and controlled by the synaptonemal complex – a conserved protein structure that assembles between homologs. The structure is built from two main elements – axes and the synaptonemal complex central region (SC-CR; Figure 1A-B). The axes are composed of cohesins and HORMA-domain proteins, which mold the chromosome into an array of loops. The SC-CR, made of coiled-coil proteins, associates with pre-assembled axes on the two homologs, placing them parallel to one another and ∼150nm apart. Synaptonemal complex assembly, or synapsis, extends localized pairing interactions to align the homologs end-to-end and intimately juxtapose homologous sequences. The synaptonemal complex also directly regulates factors that form crossover (Libuda *et al*. 2013), potentially by regulating their diffusion along chromosomes (Morgan *et al*. 2021; Zhang *et al*. 2021; Durand *et al*. 2022; Fozard *et al*. 2023).

**Figure 1:**
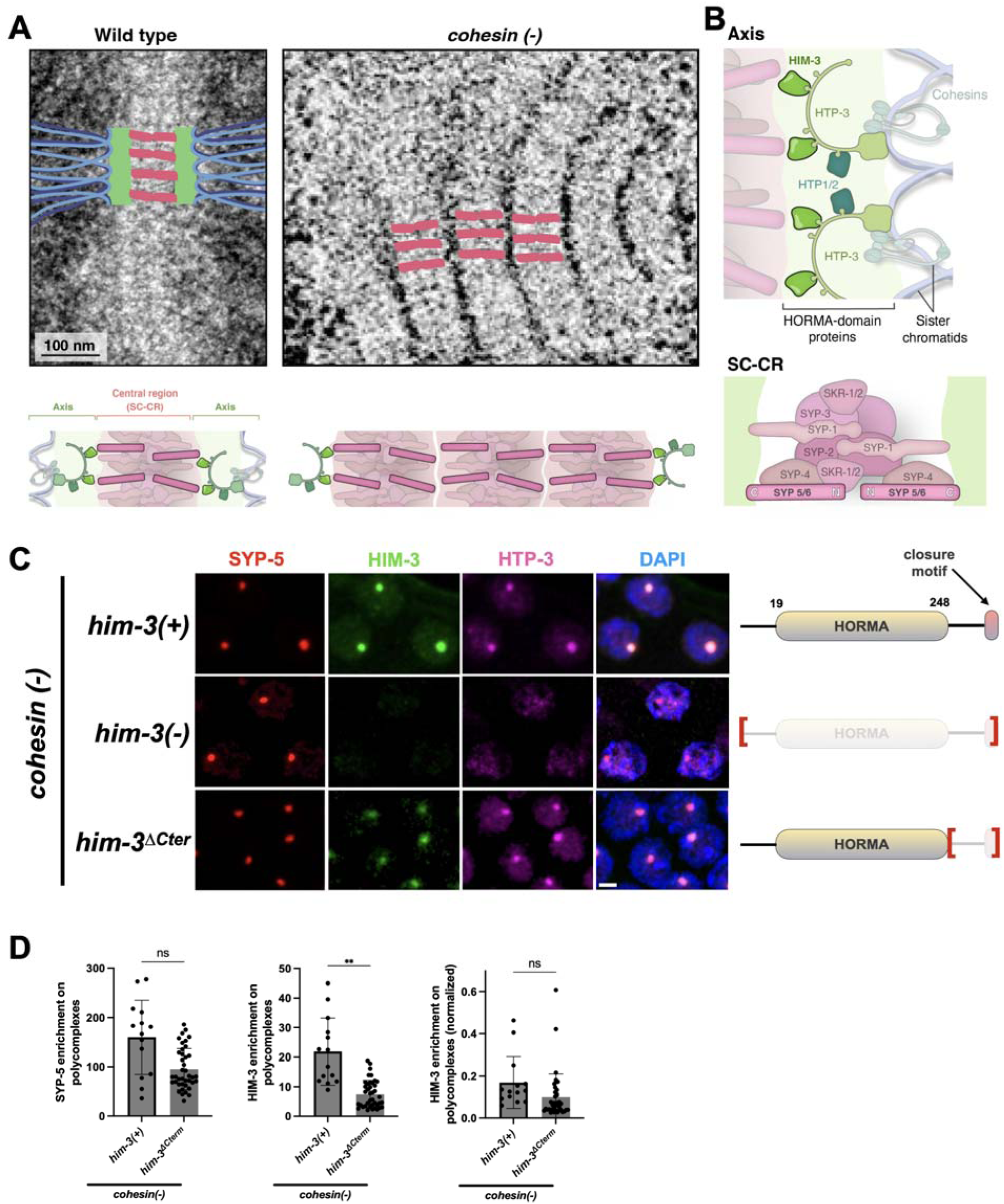
The HORMA domain of HIM-3 is required for axis interactions with the SC-CR. (A) Top left, assembled synaptonemal complex with the darkly-stained parental chromosomes to its side (left) and of the polycomplexes that form in *cohesin(-)* worms (right) as seen in negative negative-stain electron-micrographs (adapted from (Rog *et al*. 2017)). Bottom left, interpretive diagrams colored magenta for the SC-CR, green for the axes (also called lateral or axial elements) and blue for chromatin. (B) Models depicting the worm components of the axis (top) and the SC-CR (bottom). The position of each of these components within the synaptonemal complex is based on (Köhler *et al*. 2017, 2020; Hurlock *et al*. 2020; Zhang *et al*. 2020; Blundon *et al*. 2024). The pairs HTP-1/2, SYP-5/6 and SKR-1/2 are each partially redundant with each other. (C) Pachytene nuclei from worms of the indicated genotypes stained for the SC-CR component SYP-5 (red) and the axis components HIM-3 (green) and HTP-3 (magenta). The merged images on the right also show DNA (DAPI, blue). The HTP-3 antibody weakly cross-reacts with the nucleolus. Scale bar = 1 μm. Gene models of HIM-3, with the HORMA domain and the closure motif highlighted, are shown to the right. Regions deleted are denoted by red brackets. See Figure S1A for images of the gonads. (D) Quantification of the images in panel A. The enrichment at polycomplexes relative to the nucleoplasm was done using line scans. Normalized HTP-3 enrichment was calculated by dividing HTP-3 enrichment by SYP-5 enrichment.

The mechanism of synaptonemal complex assembly remains unknown. The ladder-like appearance of the SC-CR in negative-stained electron micrographs (Figure 1A), the stereotypic organization of subunits within the SC-CR (Figure 1B; (Schild-Prüfert *et al*. 2011; Schücker *et al*. 2015; Köhler *et al*. 2020)), and its assembly through processive extension (Rog and Dernburg 2015; Pollard *et al*. 2023) all contributed to the idea that assembly proceeds through zipping. This mode of assembly would be similar to active polymerization -locally consuming free energy generated by NTP hydrolysis to attach subunits at the growing end and in this way resist the restoring force of chromatin. The more recent observations of constant SC-CR subunit exchange within the synaptonemal complex and of fluid behaviors exhibited by the SC-CR suggest that it is a biomolecular condensate with liquid properties (Rog *et al*. 2017; Pattabiraman *et al*. 2017; Nadarajan *et al*. 2017; von Diezmann *et al*. 2024). The synaptonemal complex may therefore assemble by condensation of the SC-CR between parallel axes, moving chromosomes by capillary-like forces. However, available tools to reconstitute, perturb, and image the synaptonemal complex have failed to distinguish between possible assembly mechanisms. Underlying these challenges is the inability to modulate the interactions between the SC-CR and the axis, since the molecular contacts between them are not known (Gordon and Rog 2023).

Here, we identify components of the axis-SC-CR interface in the nematode *Caenorhabditis elegans*, comprising a conserved positive patch on the axis protein HIM-3 and the C-terminus of the SC-CR protein SYP-5. The positive patch on HIM-3 interacts with SYP-5 *in vivo* and *in vitro* to mediate synaptonemal complex assembly. The effects of weakened axis-SC-CR interactions on the morphology of the synaptonemal complex support SC-CR assembly through wetting. To substantiate this idea, we generated a thermodynamic model. Our model assumes no local consumption of free energy and relies on the condensation of SC-CR molecules and on surface binding of SC-CR components to the axis to account for the experimentally observed phenotypes of meiotic perturbations.

## Results

### The axis protein HIM-3 is a component of the axis-SC-CR interface

To identify the axis-SC-CR interface, we wanted to study this interface independently of other mechanisms that affect chromosome organization. We used polycomplexes: assemblies of SC-CR material that form when the SC-CR cannot load onto chromosomes. Since the stacked SC-CR lamellae in polycomplexes closely resemble the SC-CR layer that forms between the axes under physiological conditions, polycomplexes have been used to study SC-CR ultrastructure (Sym and Roeder 1995; Hughes and Hawley 2020). We used worms that lack meiotic cohesins (deletion of the meiotic kleisins *rec-8* and *coh-3/4*, designated *cohesin(-)*), which prevents axis assembly onto chromosomes. In these worms, SC-CR material forms chromatin-free polycomplexes that recruit axis components (Figures 1A, 1C and S1B; (Severson and Meyer 2014; Rog *et al*. 2017)).

First, we wanted to identify the axis components required for polycomplex-axis interactions. Out of the four meiotic HORMA proteins in worms - HTP-3, HIM-3 and HTP-1/2 - we predicted a crucial role for HIM-3 based on its proximity to the SC-CR and the increasing synapsis defects upon its gradual removal (Kim *et al*. 2015; Köhler *et al*. 2017; Gordon and Rog 2023). Upon deletion of *him-3*, the axis components HTP-3 and HTP-1/2 failed to localize to polycomplexes, as revealed by immunostaining (Figure 1C-D). This suggested that HIM-3 directly interacts with the SC-CR, whereas HTP-3 and HTP-1/2 are recruited to polycomplexes indirectly, through interactions with HIM-3 (Kim *et al*. 2014). Sequential deletions of HIM-3 regions showed that the C-terminus of HIM-3, which includes a disordered linker and a domain that interacts with other HORMA proteins (called the ‘closure motif’), plays only a minimal role in the recruitment of axis proteins to polycomplexes (Figure 1C-D). This result suggested that the SC-CR-interacting region lies in the HORMA domain of HIM-3.

The HORMA domain is a conserved fold shared among meiotic axis proteins (Ur and Corbett 2021). We examined the structures of the HORMA domains in the three meiotic axis proteins - HIM-3, HTP-3 and HTP-1/2 (Figure 2A; (Kim *et al*. 2014)) - to identify divergent surfaces that could mediate HIM-3’s specific contribution to axis-SC-CR interactions. We noticed a positively charged patch that is unique to HIM-3, containing lysines at positions 170, 171, 177 and 178 and an arginine at position 174 (Figure 2A and S2A). This positive patch is unique to HIM-3 among *C. elegans* HORMA proteins, but a similar positive patch is also present in HORMA-domain containing axis proteins in other species (Figure S2B). Specifically, HORMA axis proteins in budding yeast and animals possess a pair of positively-charged residues that align with lysines 170 and 171 on HIM-3. In addition, positively-charged residues are present in other solvent-exposed positions of the same alpha helix, such as the residues homologous to lysine 162 on HIM-3. Structural analysis confirmed that the basic residues of *S. cerevisiae* Hop1 (arginines 175 and 176) and human HORMAD1 (arginine 155 and lysine 156) occupy an equivalent position to lysines 170 and 171 on HIM-3 (Figure S2C).

**Figure 2:**
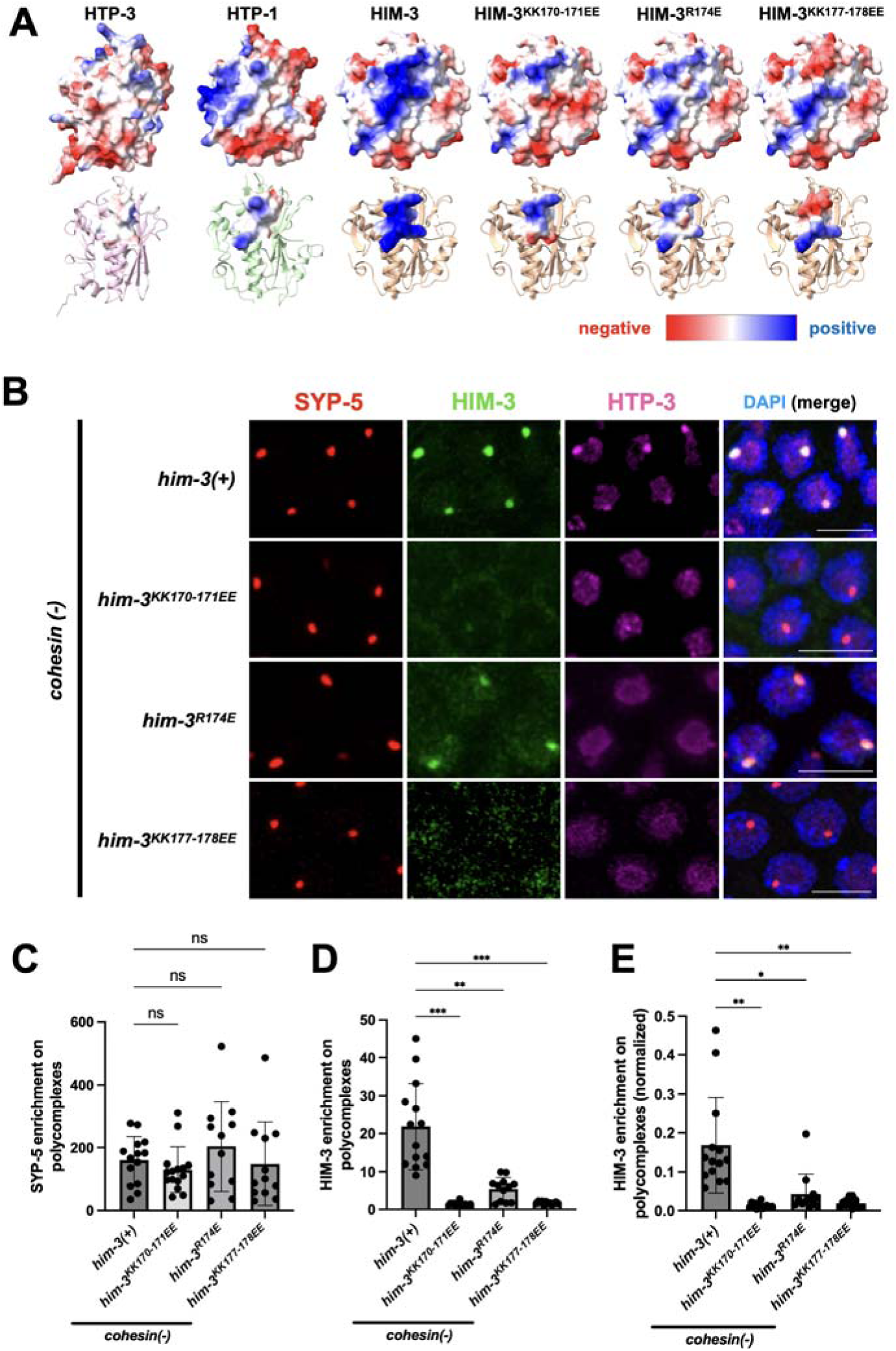
Axis interactions with the SC-CR are mediated by a positive patch on the HORMA domain of HIM-3. (A) Structural models of the meiotic HORMA proteins, with surface charge plotted in a red-blue scale. The structures of HTP-1 and HIM-3 are from (Kim *et al*. 2014). The models of HTP-3 and the three HIM-3 mutants were generated in AlphaFold (Senior *et al*. 2020). Bottom, secondary structural models, with amino acids constituting the positive patch on HIM-3 (positions 170, 171, 174, 177 and 178), and the analogous positions in HTP-1 and HTP-3 are shown as surfaces colored according to charge. (B) Pachytene nuclei from worms of the indicated genotypes stained for the SC-CR component SYP-5 (red) and the axis components HIM-3 (green) and HTP-3 (magenta). The merged images on the right also show DNA (DAPI, blue). The HTP-3 antibody weakly cross-reacts with the nucleolus. Scale bars = 1 μm. See Figure S2D for images of the gonads and Figure S3 for similar analysis in live gonads. (C-E) Quantification of the images in panel B. The enrichment at polycomplexes relative to the nucleoplasm was done using line scans. Normalized enrichment (panel E) was calculated by dividing HIM-3 enrichment by SYP-5 enrichment.

We generated several HIM-3 mutants that reversed the charge in the positive patch (Figure 2A). These mutations decreased the accumulation of HIM-3 on polycomplexes: ∼4-fold for *him-3^R174E^* and reduction to almost background level for *him-3^KK170-171EE^* and *him-3^KK177-178EE^* (Figure 2B-E; in panel E we normalized HIM-3 enrichment relative to SYP-5 enrichment). The indirect recruitment of HTP-3 to polycomplexes was also abolished. Analysis in live *cohesin(-)* gonads, using GFP-tagged HIM-3, yielded similar results (Figure S3; ∼5-fold reduction for *him-3^R174E^* versus wild-type worms). These data indicate that the positive patch on HIM-3 mediates association with SC-CR components.

### The HIM-3 positive patch is essential for synaptonemal complex assembly

To assess the contribution of the HIM-3 positive patch to synapsis, we analyzed meiosis in our HIM-3 positive patch mutants. Worms harboring *him-3^R174E^* and *him-3^KK170-171EE^* exhibited disrupted meiosis, consistent with their relative disruption of axis-SC-CR interactions. The *him-3^R174E^* worms had only 21 progeny on average, as compared with 300 progeny for wild-type worms, with 4.2% male self-progeny, indicative of mis-segregation of the *X* chromosome (Figure 3A-B; wild-type worms have 0.1% male progeny). These defects were much more severe in *him-3^KK170-171EE^* worms, which exhibited phenotypes similar to *him-3* null worms (Couteau *et al*. 2004) and were almost sterile.

**Figure 3:**
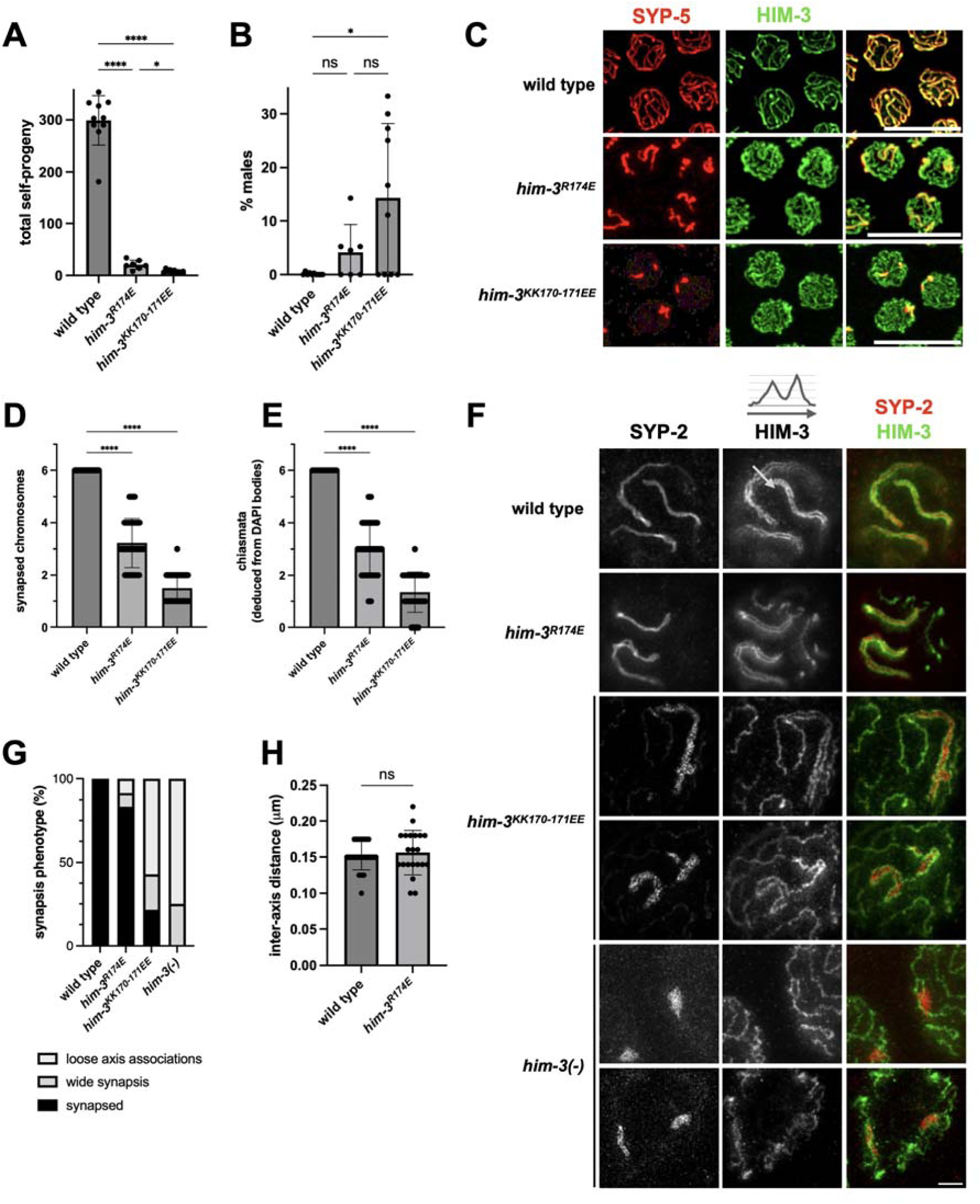
Lowering SC-CR affinity for the axes perturbs synapsis. (A) Total self-progeny from hermaphrodites of the indicated genotypes. (B) Percentage of males among self-progeny of hermaphrodites of the indicated genotypes, indicative of meiotic *X* chromosome non-disjunction. (C) Pachytene nuclei stained for the SC-CR component SYP-5 (red) and the axis component HIM-3 (green), with merged images shown on the right. Note the extensive asynapsis in the *him-3* mutants (i.e., axes lacking SC-CR staining) despite loading of the mutated HIM-3 proteins onto the axis. Scale bars = 10 μm. See Figure S4A for images of the gonads. (D) Quantification of the images in panel C, indicating a smaller number of synapsed chromosomes in *him-3* mutants. (E) Chiasmata number deduced from the number of DAPI bodies at diakinesis. Wild-type animals undergo one chiasma per chromosome, for a total of six chiasmata per nucleus. (F) STED microscopy images of pachytene nuclei stained for the SC-CR component SYP-2 (red in the merged image) and the axis component HIM-3 (green in the merged image). An example of a line scan through a synapsed chromosome is shown above the HIM-3 staining in wild-type animals. Scale bar = 1 μm. (G) Quantification of different synapsis phenotypes in STED images, as shown in panel (F). ‘Wide synapsis’ indicates parallel axes separated by more than 150nm, as shown in the top nucleus from *him-3^KK170-171EE^* animals. ‘Loose axis associations’ indicate axes wrapped around SC-CR structures, as shown in the bottom nucleus from *him-3^KK170-171EE^* animals. (H) Inter-axes distance in the indicated genotypes, measured from nuclei stained as in panel F. Distance was measured only between parallel axes that had unilamellar SYP-2 staining.

Cytological examination indicated that an average of 3.2 out of the six chromosome pairs synapsed in *him-3^R174E^* worms, and only 1.5 chromosomes synapsed in *him-3^KK170-171EE^* worms (Figure 3C-D and S4A). The synapsed chromosomes appear to form crossovers, as indicated by the correspondence between the number of synapsed chromosomes and the number of chromosomes attached through chiasmata (Figure 3E). The residual association of SC-CR material with axes in *him-3^KK170-171EE^* worms suggests that other axis components may harbor a weak affinity for the SC-CR. Consistent with this idea, chromosomes with axes that lack HIM-3 altogether are still associated with SC-CR material (Figure 3F; (Kim *et al*. 2014)).

Importantly, HIM-3 positive patch mutant proteins still loaded onto both synapsed and asynapsed chromosomes (Figure 3C). HIM-3 levels and the fraction of HIM-3 on chromosomes were also minimally affected in the mutants (Figure S4B-D). These data suggest that the surface charge alterations in *him-3* are *bona fide* separation-of-function mutations, and that the phenotypes they exhibit can be attributed to disrupted axis-SC-CR interactions.

### Disrupting axis-SC-CR interactions alters synaptonemal complex morphology

To gain better insight into the morphology of the synaptonemal complex in *him-3* mutants, we used stimulated emission-depletion super-resolution microscopy (STED). The axes in wild-type worms, and in most synapsed chromosomes in *him-3^R174E^* worms, exhibited the canonical layered ultrastructure of an assembled synaptonemal complex: they were parallel along their length, separated by ∼150nm (Figure 3F-H; (Page and Hawley 2004; Almanzar *et al*. 2023; Zickler and Kleckner 2023)). *him-3^KK170-171EE^* and *him-3(-)* chromosomes, however, were much more disorganized. Axes were often associated with each other without being parallel and even seemingly aligned axes failed to maintain a 150nm spacing (Figure 3G). In some cases, SC-CR aggregates interacted with multiple axes -a situation never observed in wild-type worms (Figure 3F-G).

Staining a protein that localizes in the middle of the SC-CR (SYP-2; Figure 1B; (Schild-Prüfert *et al*. 2011; Köhler *et al*. 2020)) revealed that many of the SC-CR structures in *him-3^KK170-171EE^* worms do not form the single thread observed in wild-type animals, implying the inter-axes space is occupied by more than a single lamella of SC-CR (Figure 3F). Instead, the SYP-2 epitope exhibited a dotty appearance with some parallel threads (Figures 3F and S5). Measurements in live worms confirmed the presence of many more SC-CR molecules per chromosome in *him-3^KK170-171EE^* worms compared with wild-type or *him-3^R174E^* worms (Figure S11F). This pattern is reminiscent of polycomplexes, which resemble stacked SC-CR lamellae, with a distance between the center of each lamella (where SYP-2 localizes) matching the width of native synaptonemal complex (Figure S5; (Rog *et al*. 2017; Hughes and Hawley 2020)).

Taken together, our analyses indicate that HIM-3-mediated axis-SC-CR interactions drive synaptonemal complex assembly. Furthermore, the altered SC-CR morphology in *him-3* mutants sheds light on the mechanism of synapsis, pointing to an interplay between axis-SC-CR interactions and self-interactions among SC-CR subunits. Below, we use this understanding to generate a thermodynamic model for synaptonemal complex assembly.

### The SC-CR protein SYP-5 interacts with the HIM-3 positive patch

To identify SC-CR components that interact with the HIM-3 positive patch, we searched the worm SC-CR subunits - SYP-1-6 and SKR-1/2 - to identify those that harbor negatively charged regions that localize near the axes. An attractive candidate was SYP-5, which has a negatively charged C-terminus that localizes near the axes and, when mutated, leads to synapsis defects (Figure 1B; (Hurlock *et al*. 2020; Zhang *et al*. 2020; Gold *et al*. 2024). (The C-terminus of SYP-1, which also localizes near the axes, is not negatively charged.)

We generated two charge-swap mutants in *syp-5* (*syp-5^5K^* and *syp-5^6K^*, mutating five and six aspartic and glutamic acids to lysines, respectively; Figure 4A). We analyzed them in the *cohesin(-) him-3^KK170-171EE^* background, hypothesizing they may restore the recruitment of axis components to polycomplexes. We found that polycomplexes in *cohesin(-) him-3^KK170-171EE^ syp-5^5K^* worms recruited significantly more HIM-3 compared to *cohesin(-) him-3^KK170-171EE^* controls (Figure 4B-E). This likely underestimates the effect of *syp-5^5K^*on axis-SC-CR interactions, since polycomplexes in this background concentrated much less SC-CR, likely due to impaired SC-CR self-interactions (Figures 4C and S11B; (Zhang *et al*. 2020)). The *syp-5^6K^* mutation further weakened SC-CR self-interactions, completely preventing polycomplex formation in *cohesin(-) him- 3^KK170-171EE^* worms and precluding assessment of its effect on axis-SC-CR interactions (Figure 4B). These data suggest that the C-terminus of SYP-5 contributes to axis-SC-CR interactions, in addition to promoting self-interactions between SC-CR subunits.

**Figure 4:**
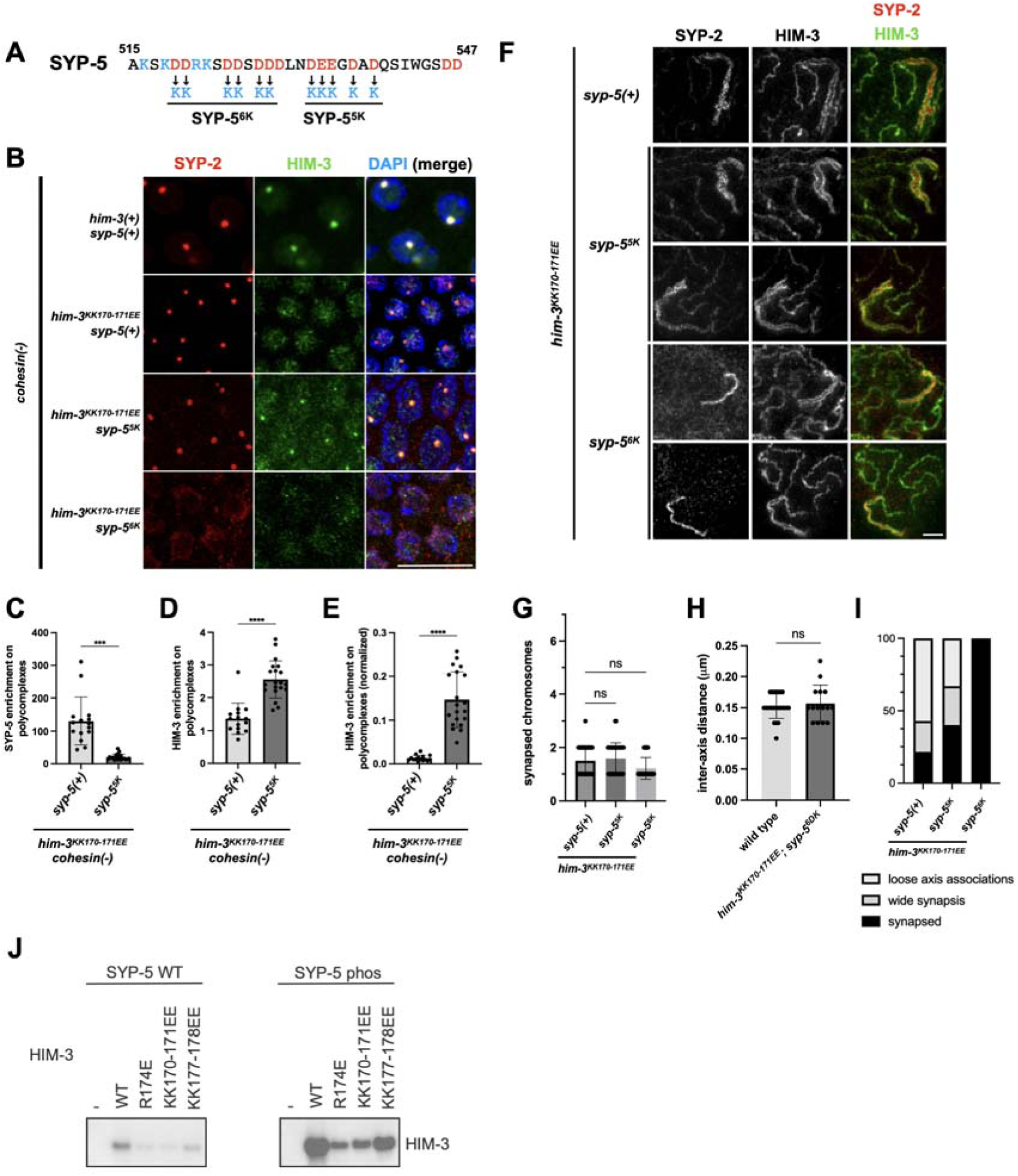
The C-terminus of SYP-5 contributes to SC-CR interactions with the axis. (A) The C-terminus of SYP-5 (amino acids 515-547), with positively-and negatively-charged residues colored in blue and red, respectively. Below, *syp-5* mutations flipping charges in the C-terminus. (B) Pachytene nuclei of the indicated genotypes stained for the SC-CR component SYP-2 (red) and the axis component HIM-3 (green). The merged images on the right also show DNA (DAPI, blue). Note that the *syp-5^6K^* mutant fails to form polycomplexes, likely due to perturbed self-interactions of the SC-CR. Scale bars = 10 μm. See Figure S6A for images of the gonads. (C-E) Quantification of the enrichment of SYP-2 and HIM-3 to polycomplexes. While the SC-CR is less enriched at polycomplexes in *syp-5^5K^*animals, these polycomplexes recruit more HIM-3. In panel E, HIM-3 enrichment is normalized to the level of SYP-2 enrichment. (F) STED microscopy images of pachytene nuclei stained for the SC-CR component SYP-2 (red in the merged image) and the axis component HIM-3 (green in the merged image). Scale bar = 1 μm. (G) The number of SC-CR structures per pachytene nuclei in the indicated genotypes. (H) Inter-axes distance in the indicated genotypes, measured from nuclei stained as in panel F. Distance was measured only for *him-3^KK170-171EE^ syp-5^6K^* mutants, where the parallel axes exhibited unilamellar SC-CR staining, and is compared to the data from Figure 3H. (I) Quantification of different synapsis phenotypes in STED images, as shown in panel F. See Figure 3G for more details. (J) Anti-HIM-3 western blot showing the interaction of purified HTP-3–HIM-3 complexes containing wild type (WT) or indicated mutants of HIM-3 with biotinylated peptides spanning residues 528-547 of SYP-5, either unmodified or phosphorylated at S541 (SYP-5 Phos). See Figure S8 for complete blots and additional controls.

When we analyzed *him-3^KK170-171EE^ syp-5^5K^* and *him-3^KK170-171EE^ syp-5^6K^* worms, we found only one or two SC-CR-associated chromosomes per nucleus, similar to *him-3^KK170-171EE^* worms (Figures 4F-G and S6B). Crucially, however, the synaptonemal complex on these synapsed chromosomes exhibited morphologies more similar to wild type. This effect was the strongest for *him-3^KK170-171EE^ syp-5^6K^* worms, where almost all the synaptonemal complexes exhibited a canonical morphology: a single SC-CR thread between the axes and an inter-axis distance of ∼150nm (Figure 4H-I).

When analyzed by themselves, both *syp-5^5K^* and *syp-5^6K^*worms exhibited defects in synaptonemal complex assembly (Figure S7A-B). These defects included the presence of asynapsed chromosomes and chromosomes that failed to form a crossover, as well as consequent defects in chromosome segregation leading to reduced progeny number and a higher prevalence of male self-progeny. Consistent with its stronger effect on synaptonemal complex morphology in *him-3^KK170-171EE^* worms, *syp-5^6K^*worms exhibited stronger defects compared with *syp-5^5K^* worms.

To confirm that HIM-3 interacts directly with the C-terminus of SYP-5, we expressed and purified a complex of HTP-3 bound to multiple HIM-3 molecules (Kim *et al*. 2014) and tested binding to a peptide spanning residues 528-547 of SYP-5. We also tested binding of the HTP-3–HIM-3 complex to a SYP-5 peptide phosphorylated at serine 541; phosphorylation of this site by polo-like kinases was recently suggested to be important for synapsis (Gold *et al*. 2024). HIM-3 strongly interacted with the phosphorylated SYP-5 peptide (Figures 4J and S8). This interaction was much weaker with unphosphorylated SYP-5 peptide, further underscoring the importance of this modification (Gold *et al*. 2024). HTP-3 alone did not interact with the SYP-5 peptides, demonstrating that HIM-3 is required for this interaction (Figure S8).

The interaction between HIM-3 and the SYP-5 peptides was dramatically reduced upon charge swap mutations on the HIM-3 positive patch. HIM-3^R174E^, HIM-3^KK170-171EE^ and HIM-3^KK177-178EE^ all interacted to a similar level with HTP-3 (Figures 4J and S8), consistent with the ability of charge swap mutants to assemble axes *in vivo* (Figures 3 and S4). However, all three mutants exhibited reduced binding to the SYP-5 peptides, demonstrating the specificity of HIM-3-SYP-5 interaction and the importance of the positive charge. The drastic *in vitro* effects of *him-3^R174E^ versus* its milder effects *in vivo* is likely due to presence of additional binding surfaces on SC-CR components that interact with arginine 174. Taken together with our *in vivo* data, we conclude that axis-SC-CR interactions are mediated by direct interaction between the positive patch on HIM-3 and the C-terminus of SYP-5 to promote synaptonemal complex assembly.

### Thermodynamic model of synaptonemal complex assembly

Our analysis of *him-3* mutants helps differentiate between different mechanisms of synaptonemal complex assembly. Zipping-based mechanisms predict that disrupting axis-SC-CR interactions will not prevent zipping *per se* but will affect the alignment of the axes (and the chromosomes) by decoupling the axes from the SC-CR. Thermodynamically-driven assembly makes a different prediction. To assemble, condensation mediated by attractive self-interactions and surface binding to the axes overcomes the entropic-driven dispersion of SC-CR components and chromosomes. These interactions together determine the morphology of the synaptonemal complex. Our observations in *him-3^KK170-171EE^* worms support thermodynamically-driven assembly: the drastically weakened axis-SC-CR interactions led to the formation of a much thicker SC-CR that failed to extend to the entire length of the chromosome (Figures 3 and S13).

To explore whether thermodynamically-driven assembly underlies synapsis, we developed a free-energy-based model. Our model incorporates the dimensions of meiotic nuclei and chromosomes in worms (Figure 5A; see Supplementary Note 1 for a full description of the model). An important quantity in our model is the condensate volume, V . We measured V for polycomplexes (∼0.05 µm^3^; Figure 5E) and found it to be somewhat smaller than the volume of the assembled SC-CR on chromosomes (∼0.1 µm^3^; Figure 5A). That is expected given the affinity between the axes and the SC-CR, allowing to recruit more SC-CR material. Since volume is not easy to measure in fluorescent images, we also used the fraction of SC-CR molecules in condensates, either polycomplexes or assembled synaptonemal complex, as a proxy for V_C_ (e.g., in Figure 5D).

**Figure 5:**
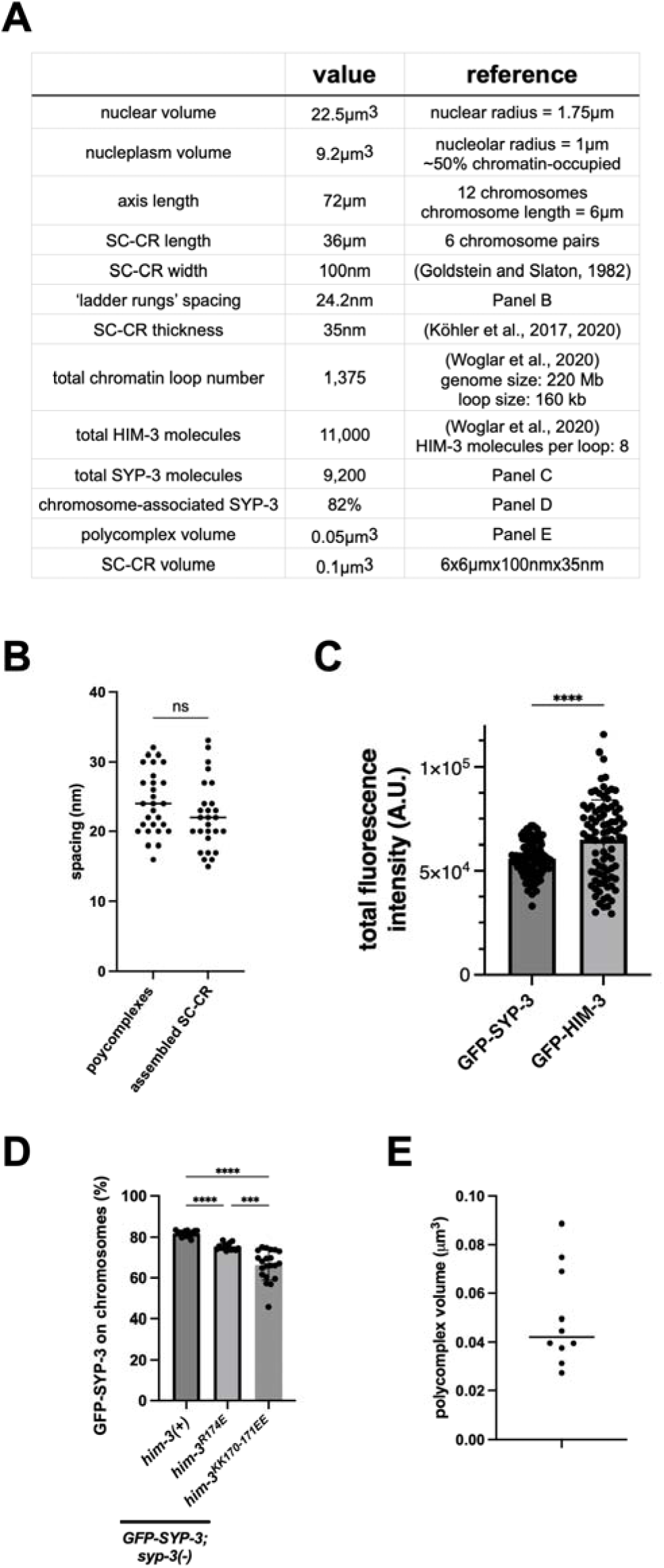
Parameters for the thermodynamic model of synaptonemal complex assembly. (A) Parameters used to model synaptonemal complex assembly. Sources: (Goldstein and Slaton 1982; Köhler *et al*. 2017, 2020; Woglar *et al*. 2020). (B) Distance between the ’ladder rungs’ in negative stain electron microscopy images from (Rog *et al*. 2017). Each point represents an individual measurement between adjacent ’rungs’. (C) Total nuclear fluorescence of GFP-HIM-3 and GFP-SYP-3 in pachytene nuclei, yielding a ratio of 1:1.2 GFP-SYP-3 to GFP-HIM-3. (D) The fraction of GFP-SYP-3 on chromosomes in animals of the indicated genotypes is significantly lower in *him-3^R174E^* and *him-3^KK170-171EE^* mutants. (E) Polycomplex volume calculated based on the dimensions of polycomplexes in negative stain electron microscopy images from (Rog *et al*. 2017). Given the mostly spherical appearance of polycomplexes, the z-dimension is assumed to be the average of the widths and height. Each point indicates a single polycomplex.

Our model includes energetic terms for two key aspects of synaptonemal complex assembly. The first is the binding of SC-CR molecules to the axis. This depends on the binding energy between SYP-5 (together with other SC-CR components) and HIM-3 (and potentially other axis components), denoted by e_SH_, as well as the number of interacting axis and SC-CR molecules. Each chromosome harbors a limited number of HIM-3 molecules (∼500), which, in turn, allow for ∼500 associated SC-CR molecules, each with binding energy e_SH_. The second free energy term incorporates the interfacial energy between the SC-CR and the nucleoplasm, which depends on attractive binding energy among SYP-5 molecules (and other SC-CR components), denoted by e_SS_, and on the minimization of the SC-CR–nucleoplasm interfacial area. The morphology of the SC-CR is therefore defined by the balance between the energetic benefit of surface binding to the axes (adsorption) and the free energy penalty of having a larger surface area for assembled synaptonemal complex threads *versus* a spherical polycomplex. While we cannot directly measure e_SS_ and e_SH_, our modeling reveals that the effects on synaptonemal complex assembly are captured by the ratio between these two entities, which we denote as 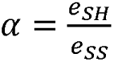.

### Synaptonemal complex assembly model recapitulates empirical observations of physiological and perturbed meiosis

The parameterized model captures multiple aspects of wild-type and mutant synapsis. First, we minimized the total free energy in the system when the condensate volume V_C_ is constant. This resulted in a monotonic relationship between a and the number of synapsed chromosomes (Figure 6A). Using this graph, the six synapsed chromosomes in wild-type worms yield a > 1.2. Similarly, the ∼3 synapsed chromosomes in *him-3^R174E^* worms translate to a = 1.0. Given the molecular nature of the mutation, this reduction in a likely reflects weaker e_SH_.

**Figure 6:**
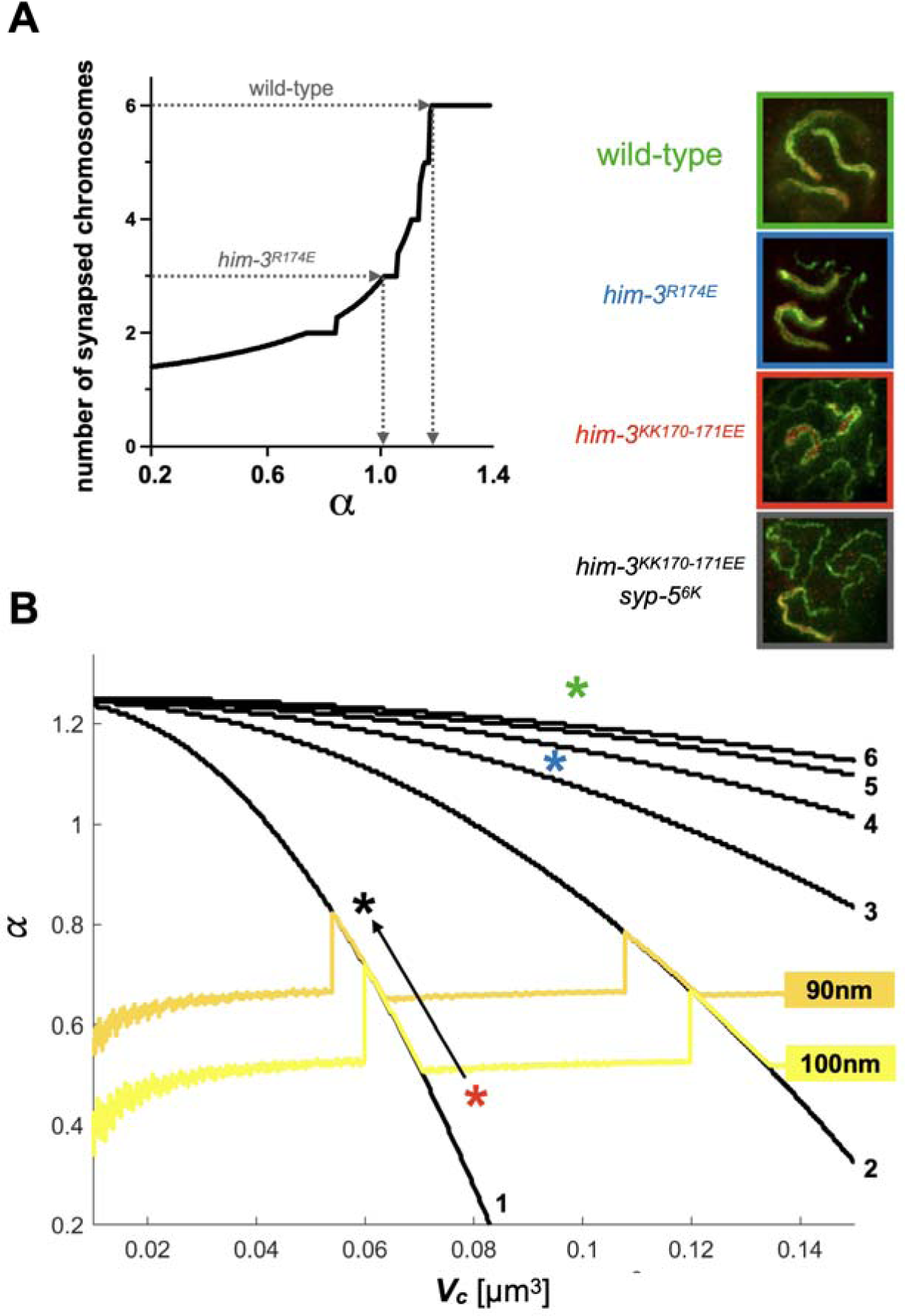
Results of the thermodynamic model of synaptonemal complex assembly. (A) Predicted number of synapsed chromosomes as a function of 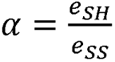. The condensate volume is held constant V*_c_* = 0.1µm^3^. Dashed arrows indicate how the number of synapsed chromosomes in wild-type and *him-3^R174E^* allows to deduce the values of a. (For simplicity, we ignore here the slight reduction [8%] in V in *him-3^R174E^* worms.) (B) Contour plot of the predicted number of synapsed chromosomes (black lines) as a function of V*_c_* and *α*. The orange and yellow lines indicate threshold SC-CR thickness of 90 and 100nm, respectively. The green, blue and red asterisks denote the position of wild-type, *him-3^R174E^* and *him-3^KK170-171EE^* worms, respectively. The black arrow and asterisk indicate the effect of combining the *syp-5* mutations with *him-3^KK170-171EE^*. Top right, example images of the mutations shown on the contour plot, with the axis stained in green and the SC-CR in red. See Supplementary Note 1 for more details.

Many of the conditions discussed here affect both a and V_C_ . To capture these complexities, we plotted the result of the model as a contour plot that links a and V_C_ to the number of synapsed chromosomes (Figure 6B; note that Figure 6A represents a simplified case, with V_C_ = 0.1). The black curves denote the minimal values of a and V_C_ that would allow the indicated number of chromosomes to synapse. On the contour plot, the wild-type and *him-3^R174E^* conditions are noted with green and blue asterisks, respectively, and *him-3^KK170-171EE^* worms, with an even lower value of a and a somewhat lower V (Figures 2 and 5D), is denoted with a red asterisk.

The contour plot also captures information about SC-CR morphology. By integrating the volume of the condensate and the number of synapsed chromosomes, we could deduce the predicted ’thickness’ of the SC-CR (i.e., the amount of material packed between the two parallel axes). Consistent with the large number of SC-CR molecules per chromosome (Figure S11F), the thickness of the SC-CR in *him-3^KK170-171EE^* worms is predicted to be >100nm (Figure 6B; thickness thresholds of 90 and 100nm are shown as orange and yellow lines, respectively). Notably, the inter-axes distance in *him-3^KK170-171EE^* worms becomes variable and the SC-CR adopting an ultrastructure resembling polycomplexes (Figures 3 and S5; (Rog *et al*. 2017; Hughes and Hawley 2020)). This suggests that only a limited amount of SC-CR material could be sandwiched between the axes while maintaining a native synaptonemal complex morphology; beyond this amount, the SC-CR forms a multi-lamellar structure.

We could similarly overlay on the contour plot the effects of other experimental perturbations. For instance, the *syp-5* mutations that partially suppress the effects of *him-3^KK170-171EE^* (Figure 3) represent diagonal upward-left vectors relative to the *him-3^KK170-171EE^* single mutant (lower V and larger a; black arrow in Figure 6B). This vector would bring the thickness of the condensate below the threshold of multi-lamellar synapsis, consistent with our empirical observations. Similarly, we model the effects of the temperature-sensitive *syp-1^K42E^* mutation, which destabilizes the SC-CR (Figure S11A; (Gordon *et al*. 2021)), and the impact of lowering the abundance of SC-CR subunits (Figure S12; see Supplementary Note 1 for full details).

Our ability to recapitulate a variety of experimental data using our free-energy-based model indicates that an active mechanism (e.g., polymerization) need not be invoked in the assembly of the synaptonemal complex. Instead, our model indicates that wetting of the axes by the SC-CR can confer selective assembly of the synaptonemal complex between homologs. We conclude that the dramatic chromosome reorganization necessary for chromosome alignment is driven by a mechanism that does not require any additional energy input beyond thermodynamics.

## Discussion

In this study, we identified a molecular contact between the axis and the SC-CR, which allowed us to explore the mechanism of synaptonemal complex assembly. Molecular genetic analysis combined with *in vivo* and *in vitro* experiments revealed an electrostatic interface between a positive patch on the HIM-3 HORMA domain and the negatively charged C-terminus of SYP-5. The residual SC-CR-axis association in worms lacking HIM-3 altogether (Figure 3F) suggests that the HIM-3-SYP-5 interaction acts together with additional contacts to form the axis-SC-CR interface.

The HIM-3 positive patch is conserved in divergent eukaryotic lineages (Figure S2B-C). Also conserved are the dimensions and ultrastructure of the synaptonemal complex (Page and Hawley 2004; Zickler and Kleckner 2023), and the SC-CR’s dynamic behaviors (Rog *et al*. 2017) and its ability to form polycomplexes (Hughes and Hawley 2020). These observations suggest that the mechanism of synaptonemal complex assembly -wetting of axes by the SC-CR - is likely to be conserved as well. Interestingly, the primary sequence of SC-CR subunits is highly divergent within and between lineages (Kursel *et al*. 2021), suggesting that sequence alignment by itself might not be sufficient to reveal the SC-CR side of the interface. Nevertheless, our observations suggest that negatively charged regions in SC-CR components could carry an analogous functional role to the C-terminus of SYP-5.

The ability to experimentally modulate the affinity between the SC-CR and the axis allowed us to test mechanisms of synaptonemal complex assembly. The liquid-like properties of the SC-CR, as demonstrated by the dynamic exchange of subunits and an ability to form droplet-like polycomplexes, led us to hypothesize that it assembles by wetting two HIM-3-coated axes. Wetting, which relies on binding (adsorption) to the axes and self-interactions between SC-CR subunits (condensation), allows the concomitant spread to the entire chromosome and the generation of adhesive forces between the homologous chromosomes. Supporting this idea, a thermodynamic model that assumes only self-interactions between SC-CR subunits and binding interactions between the SC-CR and the axis recapitulated the phenotypes of weakening axis-SC-CR and intra-SC-CR interactions (Figures 5D, 6, S10 and S11) and of lowering SC-CR levels (Figure S12).

Our synaptonemal complex assembly model provides an elegant explanation for the association of SC-CR exclusively with paired axes (MacQueen *et al*. 2005). While the SC-CR has an affinity for the axes, binding to unpaired axes provides a small energetic advantage compared with SC-CR condensation. Stable association with axes only occurs in the context of a fully assembled synaptonemal complex, where SC-CR subunits form a condensate that wets the axes. Weakening SC-CR self-association could expose the tendency of SC-CR molecules to bind unpaired axes. Indeed, two independent SC-CR mutations that weaken intra-SC-CR associations also lead to SC-CR association with unpaired axes (the mutations are the aforementioned *syp-1^K42E^* and *syp-3(me42)*; Figure S11A; (Smolikov *et al*. 2007; Rog *et al*. 2017; Gordon *et al*. 2021)).

Severe perturbations of axis-SC-CR interactions (*him-3^KK170-171EE^* or *him-3(-)*) led to the formation of large SC-CR aggregates within the axes – >400nm between the axes and too far apart to be spanned by a single SC-CR lamella (∼150nm; Figure 3F). The potential to form such a structure suggests that the wild-type scenario – where unilamellar SC-CR coats the axes from end to end – reflects a tightly regulated balance between axis-SC-CR binding and the interfacial tension of SC-CR condensates. In addition to enabling end-to-end synapsis of parental chromosomes, such a balance could also counter the thermodynamic drive of liquids to minimize surface tension (e.g., through the process of Ostwald ripening; (Gouveia *et al*. 2022)). Axis wetting therefore underlies persistent and complete synapsis – the maintenance of independent SC-CR compartments, one on each chromosome – during the many hours in which the synaptonemal complex remains assembled.

A unilamellar SC-CR has crucial functional implications. Complete synapsis ensures two fundamental characteristics of meiotic crossovers: 1) all chromosomes undergo at least one crossover and 2) crossovers only occur between homologous chromosomes. The specter of multiple axes interacting with large SC-CR aggregates (Figure 3F) is likely to prevent synapsis of all chromosomes by sequestering SC-CR material. It could also allow ectopic exchanges between nonhomologous chromosomes and, consequently, karyotype aberrations and aneuploidy. The limited surface area of a unilamellar SC-CR, together with repulsive forces between chromatin masses (Marko and Siggia 1997), could limit the number of interacting axes to no more than two. Such a mechanism to prevent multi-chromosome associations can help explain the evolutionary conservation of the synaptonemal complex, which exhibits only minor ultrastructural variations between species with order-of-magnitude differences in genome size and chromosome number (Page and Hawley 2004; Zickler and Kleckner 2023).

Our thermodynamic model groups together the distinctive affinities that drive SC-CR self-interactions: stacking of SC-CR subunits and the lateral attachments between SC-CR lamellae. The spherical morphology of stacked ladder-like lamellae in polycomplexes suggests a balance with the anisotropic elements (ladder-like assembly), and the potentially isotropic attractive interactions among SC-CR proteins. This spherical morphology is distinct from mitotic spindles (Oriola *et al*. 2020) but more akin to drops of fragmented amyloid fibrils in yeast (Tyedmers *et al*. 2010). The non-spherical polycomplexes that form in some organisms (Hughes and Hawley 2020) and in certain mutant backgrounds (e.g., (Gordon *et al*. 2021)) provide an opportunity for future studies of the balance between stacking and lateral interactions.

Cell biologists have identified numerous supramolecular assemblies in the nucleus (Sabari *et al*. 2020). Many of these structures have been suggested to exert force and movement on the genome in order to organize it and thus tightly regulate biological processes ranging from transcription to genome maintenance. The *in vitro* and *in vivo* material properties of many such structures have been a focus of recent probing. However, only rarely has it been shown that a specific material state of a supramolecular assembly (e.g., a liquid) underlies the nuclear-scale maneuvering of chromosomes in the nucleus (Gouveia *et al*. 2022; Chung and Tu 2023). Synaptonemal complex assembly through wetting demonstrates that the liquid properties of the SC-CR underlie a core component of meiosis – the large-scale chromosome reorganization that brings homologous chromosomes together.

## Supporting information

Data for Model Figure 2A

Data for Model Figure 2B

Data for Model Figure 3

## Acknowledgments

We would like to thank all members of the Rog lab and Martin Horvath for discussions and advice; Yumi Kim for discussing data prior to publication and for antibodies; Amy MacQueen for discussion and advice; Amy Strom and Erik Jorgensen for critical reading of this manuscript; Sara Nakielny for editorial work; Maria Diaz de la Loza for scientific illustrations; and Abby Dernburg for antibodies. Some worm strains were provided by the Caenorhabditis Genetics Center, which is funded by NIH Office of Research Infrastructure Programs (P40 OD010440). We acknowledge the HSC Imaging Core for the use of the STED microscope. Work in the Rog lab is funded by R35GM128804 grant from NIGMS; work in the Corbett lab is funded by R35GM144121.

## Author contributions

SGG carried out all worm experiments. AAR carried out the pull-down experiments. SGG and YG carried out phylogenetic analysis and structural predictions. CFL developed the thermodynamic model. OR and KDC supervised the experimental work. SGG, CFL, KDC and OR wrote the paper, with input from all authors.

## Supplemental information

Document S1. Figures S1–S13

Document S2. Supplemental Note 1

Table S1. Excel file containing data related to Model Figure 2A in Supplemental Note 1

Table S2. Excel file containing data related to Model Figure 2B in Supplemental Note 1

Table S3. Excel file containing data related to Model Figure 3 in Supplemental Note 1

## Materials and Methods

### Worm strains and CRISPR

Worms were grown under standard conditions (Brenner 1974). Unless otherwise noted, all worms were grown at 20°C. All strains used in this study are listed in Table S1. CRISPR was performed as previously described (Gordon *et al*. 2021), with guide RNA and repair templates listed in Table S2. All new alleles were confirmed by Sanger sequencing.

### Structural models of HORMA domain-containing proteins

PDB files for HTP-1, HIM-3 (Kim *et al*. 2014), and *H. sapiens* HORMAD1 (Wang *et al*. 2023) were downloaded from the RCSB Protein Databank (https://www.rcsb.org/) and uploaded into ChimeraX (Meng *et al*. 2023). Electrostatic models of surface charge were created with the Surfaces tab on ChimeraX. Models of *him-3* charge swap mutants were created using the Rotamers tab and changing the specified amino acids to aspartic acid residues with the ’best predicted’ position. A predicted structure of the HORMA domain of HTP-3 was generated in AlphaFold (Senior *et al*. 2020) without the C-terminal tail. The best-predicted structure was used. A predicted structure of *S. cerevisiae* Hop1 (Uniprot ID P20050) was downloaded from the AlphaFold Protein Structure Database (https://alphafold.ebi.ac.uk) (Varadi *et al*. 2024).

### Immunofluorescence and fluorescence measurements on polycomplexes

Immunofluorescence was performed as described in (Gordon *et al*. 2021). Images were acquired with a Zeiss LSM880 microscope equipped with an AiryScan and a x63 1.4NA Oil objective. The laser powers were kept the same at 1.5% 633nm, 0.3% 561nm, 2.2% 488nm and 4.5% 405nm. The antibodies used were guinea pig anti-HTP-3 (MacQueen *et al*. 2005), rabbit anti-SYP-5 (Hurlock *et al*. 2020), chicken anti-HIM-3 (Hurlock *et al*. 2020), and rabbit anti-SYP-2 (Colaiácovo *et al*. 2003), with appropriate secondary antibodies (Jackson ImmunoResearch). Line scans were analyzed in ZEN Blue 3.0 (Zeiss) on a single z-slice where the polycomplex has the highest fluorescence. The average fluorescence inside the polycomplex and in the nucleoplasm (outside the polycomplex) were used to determine enrichment on polycomplexes. To normalize, the enrichment of the axis component was divided by the enrichment of the SC-CR component.

### Meiotic phenotypes

Progeny and male counts were performed as in (Gordon *et al*. 2021). Synapsed chromosomes were counted in maximum-intensity projection images of gonads stained for an SC-CR component (SYP-2 or SYP-5). Chiasmata were counted as in (Gordon *et al*. 2021). Synapsis phenotypes were determined on STED images and were confirmed with line scans to determine that the inter-axis distance was greater than 150nm.

### STED imaging

Immunofluorescence slides were made as above, with the following modifications. We used rabbit anti-SYP-5 (Hurlock *et al*. 2020), rabbit anti-SYP-2 (Hurlock *et al*. 2020) and guinea pig anti-HTP-3 primary antibodies, and STAR RED anti-rabbit (Abberoir; 1:200) and Alexa fluor 594 anti-guinea pig (Jackson ImmunoResearch; 1:200) as secondary antibodies. We used liquid mount (Abberoir) as a mounting media. Imaging on STEDYCON was done as in (Almanzar *et al*. 2023). Line scans were used to determine the distance between axes, as described in (Almanzar *et al*. 2023).

### Live gonad imaging

Live imaging of gonads was performed essentially as described in (von Diezmann and Rog 2021). Briefly, 2% agarose pads soaked with embryonic culture medium (ECM; 84% Leibowitz L-15 without phenol red, 9.3% fetal bovine serum, 0.01% levamisole and 2 mM EGTA) for ∼20 minutes. Worms were dissected in 20µL ECM supplemented with Hoechst 33342 (1:200). The slides were sealed with VALAP and imaged using 4% 488 laser power 4.5% 405 laser power. Images were processed using Imaris 10.0 (Bitplane). 5 nuclei from each gonad were cropped and a mask for the 488 channel was made. The mask was applied using the default setting, but was manually adjusted as appropriate, particularly in some genotypes (*syp-3* RNAi and *htp-3(-)*).

### RNAi

RNAi was performed as described in (Libuda *et al*. 2013). Briefly, *syp-3* (F39H2.4) and RNAi control (pL4440) plasmids from the Ahringer laboratory RNAi library (Kamath *et al*. 2003) were grown overnight in LB+carbenicillin at 37°C, spread on RNAi plates (NGM+carbenicillin+IPTG) and incubated overnight at 37°C. L4 worms were placed on RNAi plates and grown for 24 hours at 20°C. Live gonads were imaged as described above.

### CRISPR

CRISPR/Cas9 injections were performed essentially as described in (Gordon *et al*. 2021), with the templates and guides listed in Table S1. Correct repair was confirmed by Sanger sequencing.

### Protein Purification

Codon-optimized genes encoding His_6_-HTP-3 residues 2-739 and HIM-3 residues 1-291 were cloned as a polycistronic cassette into a single vector (UCB Macrolab 2B-T; Addgene #29666), resulting in an N-terminal His_6_-tag fused to HTP-3 (Kim *et al*. 2014). For expression of HTP-3 or HIM-3 alone, single genes were cloned into the same vector. Expression vectors were transformed into *Escherichia coli* Rosetta pLysS (EMD Millipore), grown at 37°C to an OD_600_ of 0.5, then expression was induced with addition of 0.2 mM IPTG, growth temperature adjusted to 20°C, and growth continued for 16 hours. Cells were harvested by centrifugation, resuspended in lysis buffer (20 mM HEPES pH 7, 300 mM NaCl, 10% glycerol, 5 mM MgCl_2_, 5 mM imidazole, and 5 mM β-ME (β-mercaptoethanol)) and lysed in a sonicator (Branson Sonifier), then the lysate was clarified by centrifugation. Expressed proteins were purified from clarified lysates via affinity chromatography (Ni-NTA Superflow resin; Qiagen) in lysis buffer. Lysate was incubated with nickel resin for 30 minutes and then washed with lysis buffer with 20 mM imidazole. Protein was eluted in lysis buffer with 500 mM imidazole. Protein fractions were then pooled and diluted to 100 mM NaCl in 20 mM HEPES pH 7, 10% glycerol, 5 mM β-ME and loaded onto an anion exchange column (HiTrap Q HP; Cytiva). Protein was eluted with a gradient from 100 mM NaCl to 1 M NaCl, and fractions containing the desired proteins were pooled, concentrated in a centrifugal concentrator (Amicon Ultra; EMD Millipore) and loaded onto a size exclusion column (Superdex 200 Increase 10/300 GL; Cytiva) in buffer containing 20 mM HEPES pH 7, 300 mM NaCl, 10% glycerol, and 1 mM DTT. Fractions containing the desired proteins were pooled, concentrated, and stored at -80°C until needed.

### Pulldowns of HIM-3 and SYP-5

SYP-5 peptides spanning residues 528-547 (sequence: DDDLNDEEGDADQSIWGSDD) with S541 phosphorylated and unphosphorylated were purchased from Biomatik with an N-terminal biotin label. Purified HTP-3–HIM-3 was incubated with SYP-5 phosphorylated and unphosphorylated peptides (10 μg each, 25 μL each) in pulldown buffer 20 mM Tris-HCl pH 8, 150 mM NaCl, 5% glycerol, 0.01% NP-40, and 0.5 mM βME for 30 minutes at room temperature with rotation. Magnetic streptavidin resin (streptavidin beads; Vazyme) was added to protein mixture and incubated at room temperature for an additional 15 minutes with rotation. After incubation, beads were washed with 1 mL of pulldown buffer and incubated at room temperature with rotation for 5 minutes. Reaction mixture was then incubated on a magnetic strip and supernatant was removed. Pulldown buffer was added and cycles of incubating beads on magnetic strip and removing supernatant were followed until three washes were completed. Streptavidin beads were resuspended in 20 μL of 2X SDS loading buffer with 25 mM biotin and boiled. Samples were loaded onto SDS-page gel for protein identification followed by western blotting. For western blot, proteins were transferred to PVDF membrane (Bio-Rad Trans-Blot Turbo) then blocked with 3% nonfat dry milk and blotted. Blots were imaged with Bio-Rad ChemiDoc system using filters to image horseradish peroxidase (HRP) activity. Antibodies used were chicken anti-HIM-3 primary antibody (gift from Yumi Kim) at 1:5,000 dilution and donkey anti-chicken HRP conjugated secondary antibody (Jackson ImmunoResearch Laboratories Inc. #703-035-155) at 1:30,000 dilution.

### Statistical analysis

All statistical analysis was done in Prism 10.0 (GraphPad).

---

**Table S1:**
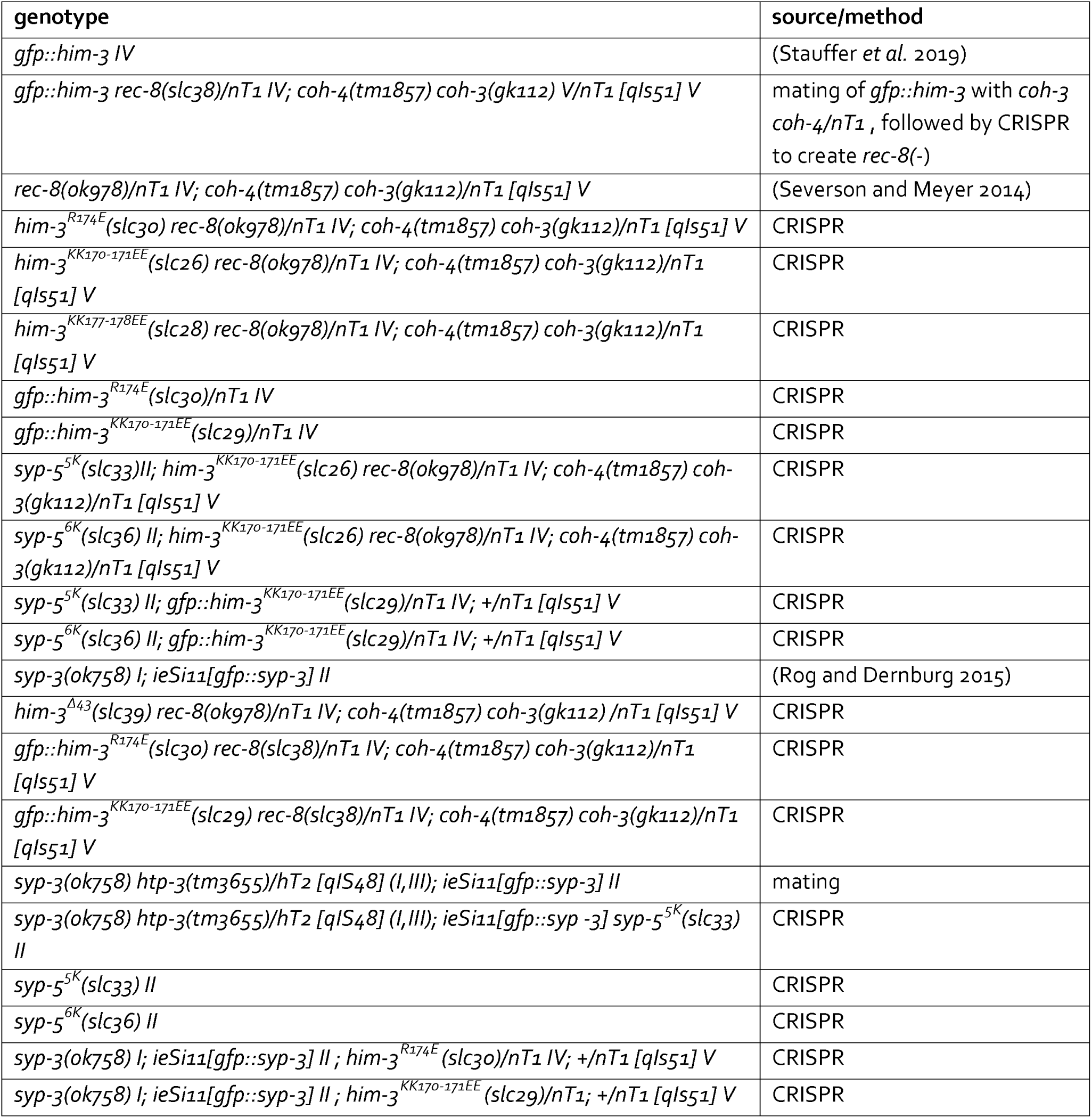
Strains used in this work.

**Table S2:**
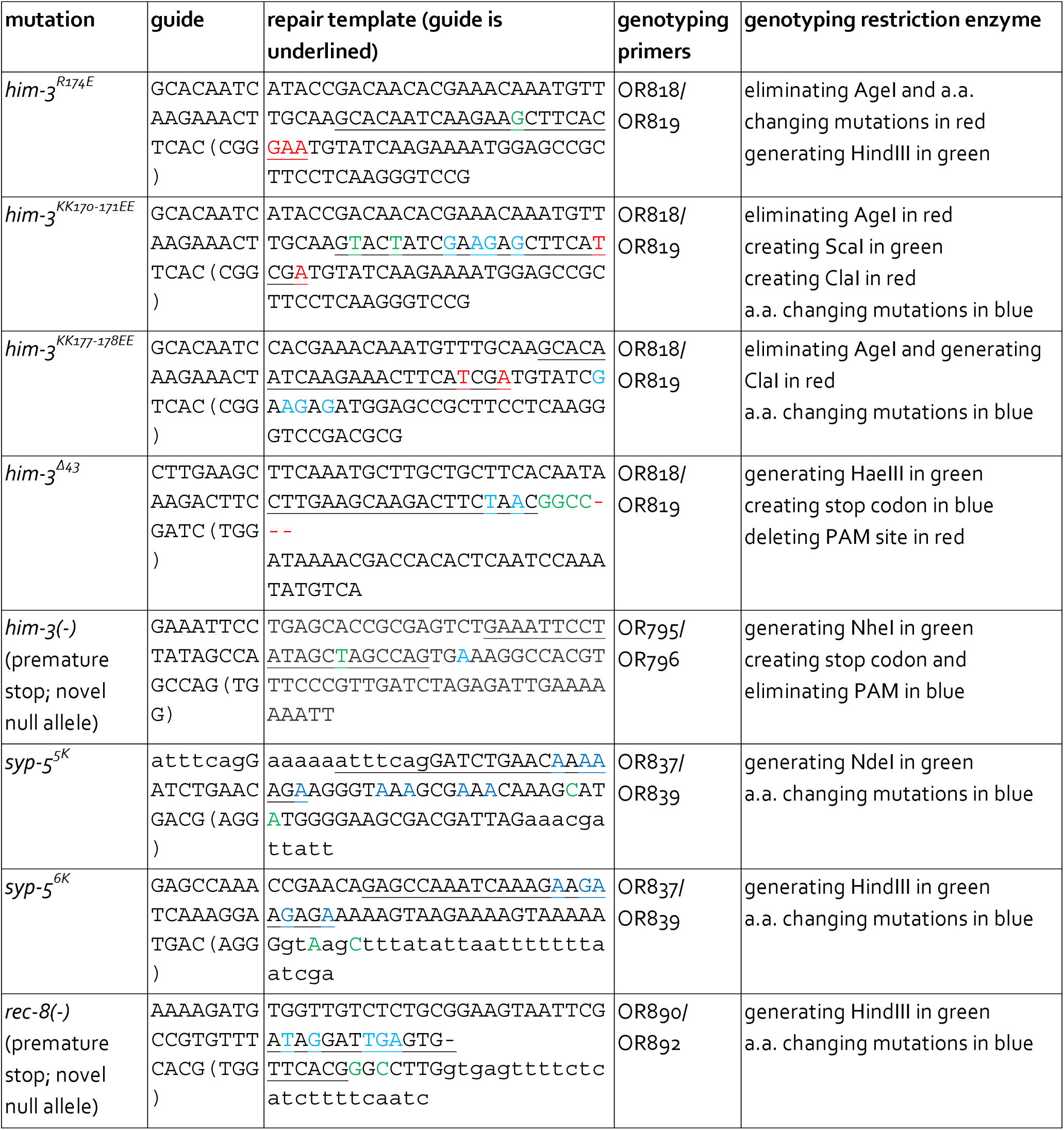
Details on CRISPR-generated alleles.

