## Supplementary Figures for "The synaptonemal complex aligns meiotic chromosomes by wetting"

**Supplemental Figures**

### Figure S1

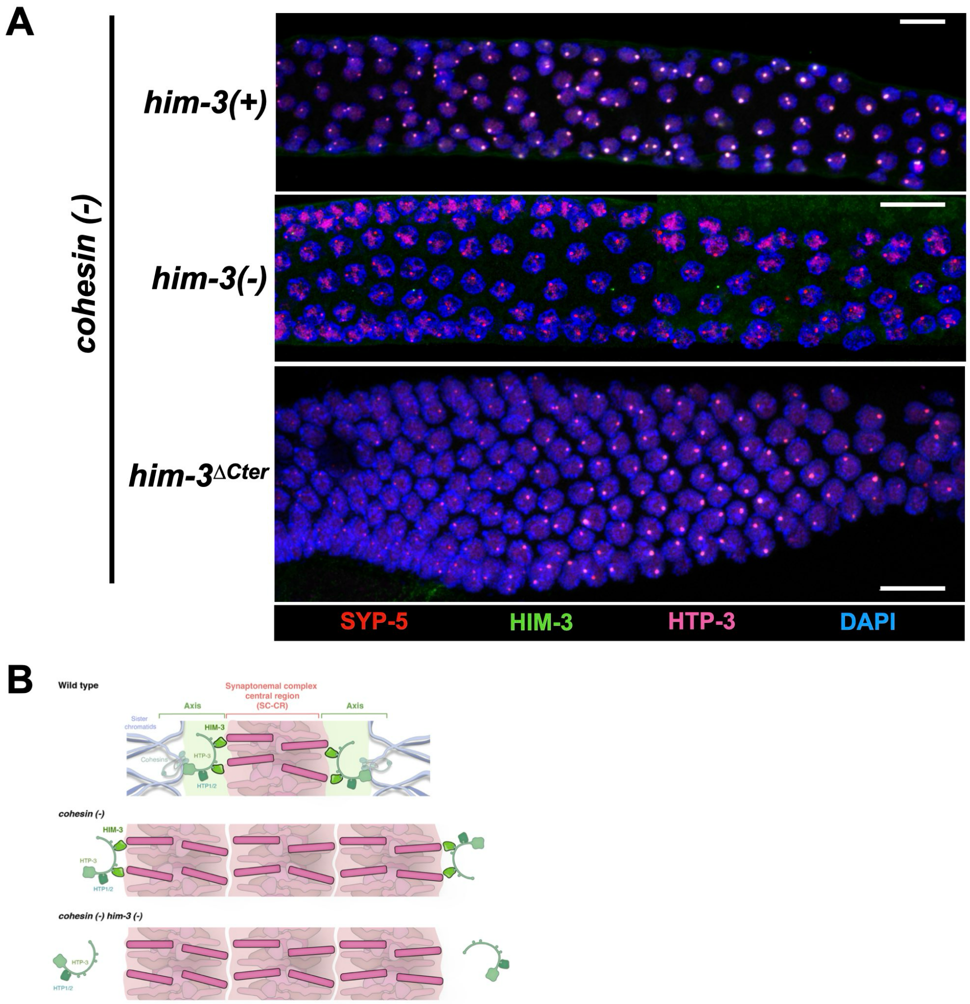


**Figure S1: Additional data relating to Figure 1**

(A) The pachytene region of gonads from worms of the indicated genotypes, corresponding to Figure 1C. Scale bars = 10 μm. (B) Graphical summary of the results in Figure 1. The polycomplexes that form in the absence of cohesins recruit the HORMA-domain-containing axis proteins. Bottom, in *him-3(-)* worms, the other axis proteins do not localize to polycomplexes.

### Figure S2

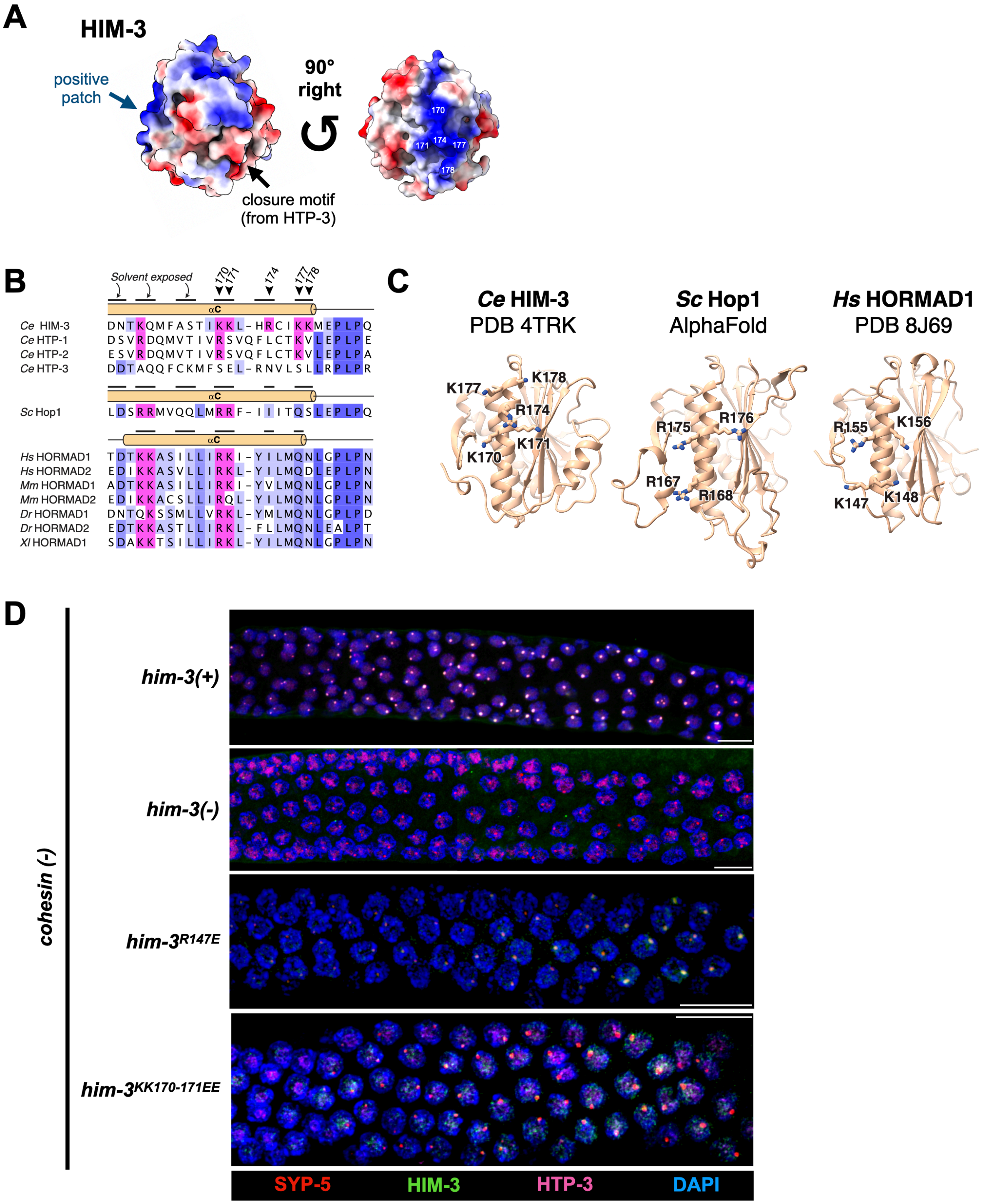
A
HIM-3
positive patch
closure motif (from HTP-3)
90° right
170
171
174
177
178
B
Solvent exposed
170
171
174
177
178
αC
Ce HIM-3 DNTKQMFASTIKKL-HRCIKKMEPLPQ
Ce HTP-1 DSVRDQMVTIVRSVQFLCTKVLEPLPE
Ce HTP-2 ESVRDQMVTIVRSVQFLCTKVLEPLPA
Ce HTP-3 DDTAQQFCKMFSEL-RNVLSLLRPLPR
Sc Hop1 LDSRRMVQQLMRRF-IIITQSLEPLPQ
Hs HORMAD1 TDTKKASILLIRKI-YILMQNLGPLPN
Hs HORMAD2 EDIKKASVLLIRKL-YILMQDLEPLPN
Mm HORMAD1 ADTKKASILLIRKI-YVLMQNLGPLPN
Mm HORMAD2 EDIKKACSLLIRQL-YILMQNLGPLPN
Dr HORMAD1 DNTQKSSMLLVRKL-YMLMQNLGPLPD
Dr HORMAD2 EDTKKASTLLIRKL-FLLMQNLEALPT
Xl HORMAD1 SDAKKTSILLIRKL-YILMQNLGPLPN
C
Ce HIM-3
PDB 4TRK
K177
K178
R174
K171
K170
Sc Hop1
AlphaFold
R175
R176
R167
R168
Hs HORMAD1
PDB 8J69
R155
K156
K147
K148
D
cohesin (-)
him-3(+)
him-3(-)
him-3R147E
him-3KK170-171EE
SYP-5
HIM-3
HTP-3
DAPI

**Figure S2: Additional data relating to Figure 2**

(A) The positive patch on HIM-3 is located away from the closure motif binding pocket. Space-filling model of HIM-3 colored according to charge, with the closure motif from HTP-3 (Kim *et al.* 2014). The positive patch is located on the other side of the HIM-3 HORMA domain. (B) Sequence alignment of the $\alpha$C helix of the four *C. elegans* (*Ce*) HORMA proteins (top), *S. cerevisiae* (*Sc*) Hop1 (middle), and other animal HORMA proteins (*Hs*: *Homo sapiens*; *Mm*: *Mus musculus*; *Dr*: *Danio rerio*; *Xl*: *Xenopus laevis*; bottom). Conservation is indicated by blue shading, with conserved positive patch residues highlighted in pink. Solvent exposed residues on the $\alpha$C helix are marked by black lines for *C. elegans* HIM-3, *S. cerevisiae* Hop1, and *H. sapiens* HORMAD1. (C) Aligned HORMA domain structures for *C. elegans* HIM-3 (PDB ID 4TRK; (Kim *et al.* 2014)), *S. cerevisiae* Hop1 (AlphaFold 2 model; (Varadi *et al.* 2024)), and *H. sapiens* HORMAD1 (PDB ID 8J69; (Wang *et al.* 2023)). Conserved positive patch residues on the $\alpha$C helix are shown as sticks and labeled. (D) The pachytene region of gonads from worms of the indicated genotypes, corresponding to Figure 2B. Scale bars = 10 μm.

### Figure S3

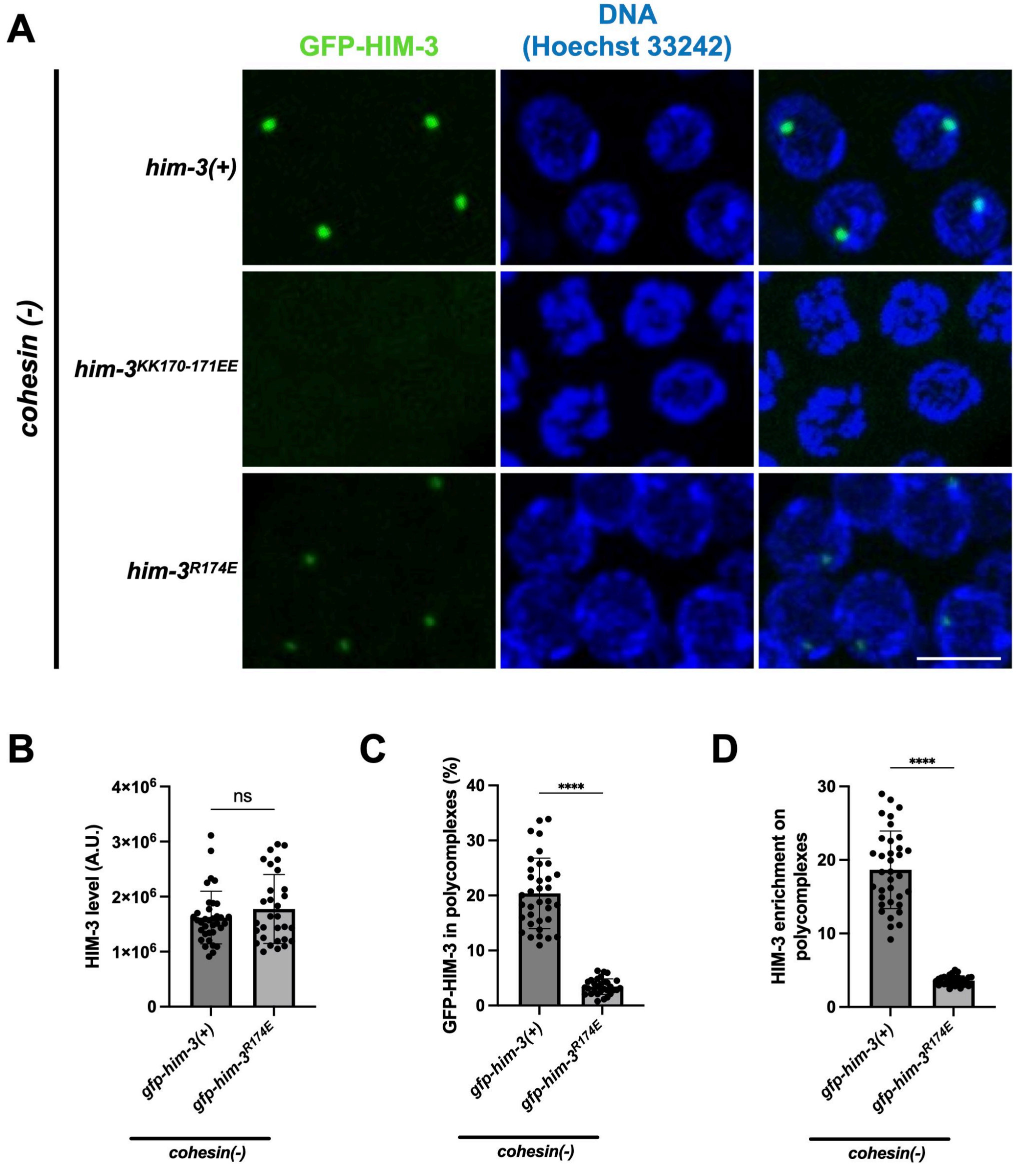


**Figure S3: HIM-3 interactions with polycomplexes measured in live gonads, relating to Figure 2**

(A) Nuclei from worms of the indicated genotypes, where HIM-3 is tagged with GFP (green). Chromatin is stained with Hoechst 33242 (blue). Similar exposure conditions were used in all cases. (B-D) Quantifications of images in panel A. GFP-HIM-3$^{KK170\text{-}171EE}$ images were not quantified since the polycomplexes could not be identified. (B) Total GFP-HIM-3 nuclear fluorescence. (C) Fraction of GFP-HIM-3 on polycomplexes. (D) Enrichment of GFP-HIM-3 on polycomplexes versus the nucleoplasm.

### Figure S4

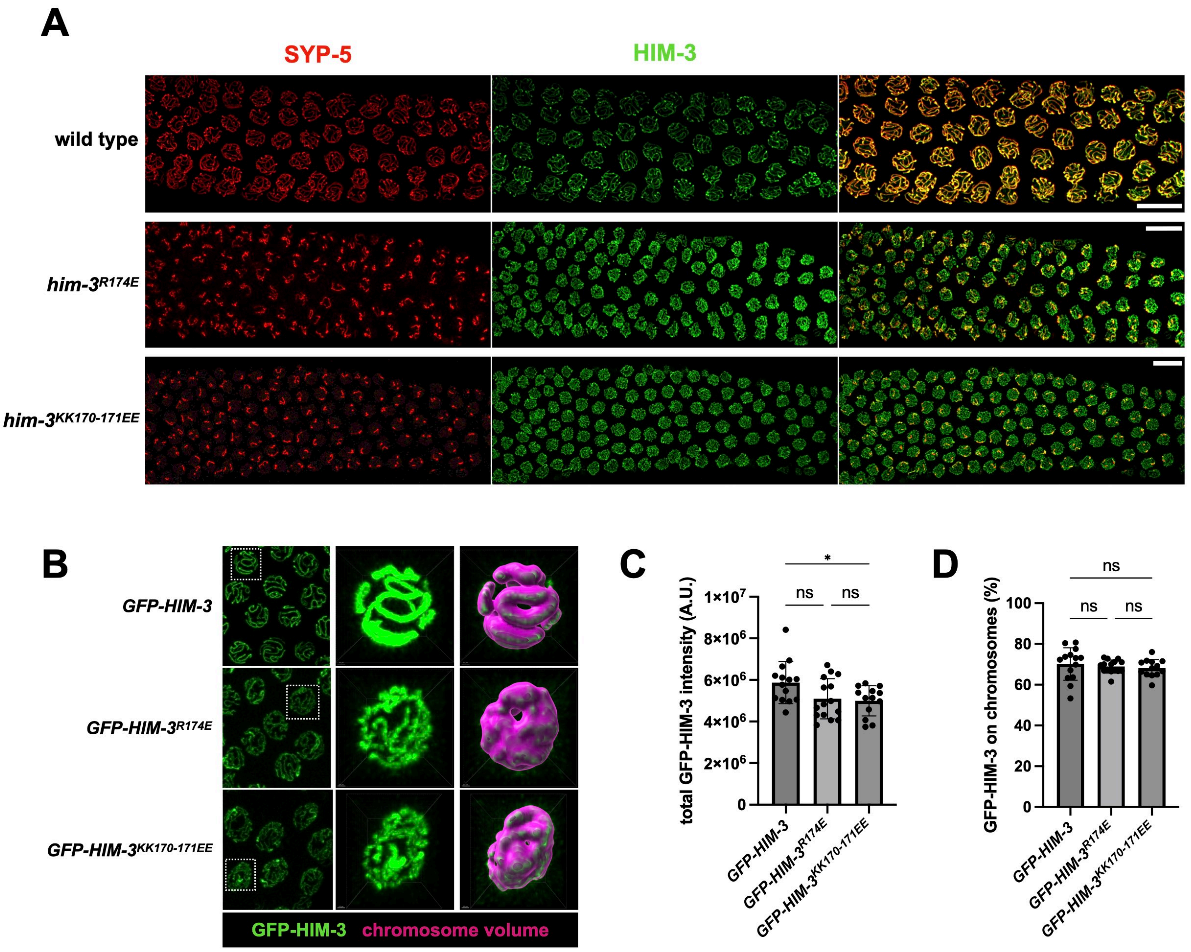


**Figure S4: Additional characterization of *him-3* mutants, relating to Figure 3**

(A) The pachytene region of gonads from worms of the indicated genotypes, corresponding to Figure 3C, stained for the SC-CR component SYP-5 (red) and the axis component HIM-3 (green), with merged images shown on the right. Scale bars = 10 µm. (B-D) Quantification of GFP-HIM-3 abundance and localization. (B) Pachytene nuclei from live worms expressing GFP-HIM-3. The middle column zooms in on a single nucleus. Right, a surface rendering of the chromosomes used to calculate the chromosome-associate fraction of GFP-HIM-3. (C) Quantification of the total amount of GFP-HIM-3 in the indicated genotypes. The minor reduction in total fluorescence in *him-3$^{KK170\text{-}171EE}$* mutants, while statistically significant, is likely not to have biological importance. (D) The fraction of GFP-HIM-3 (and *him-3* mutants) on chromosomes. Notably, *him-3* positive patch mutations do not significantly alter HIM-3 recruitment to chromosomes.

### Figure S5

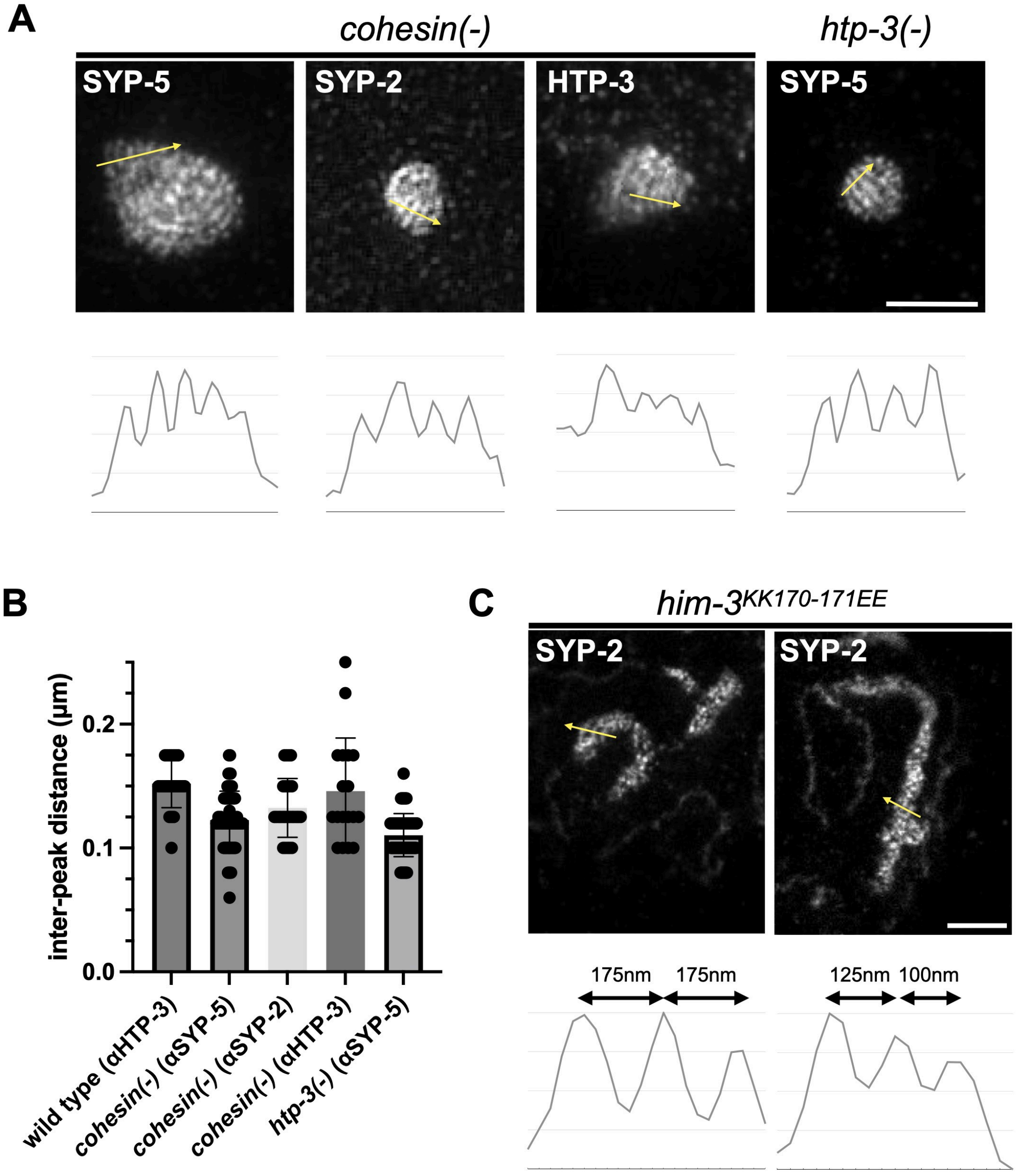


**Figure S5: SC-CR staining in *him-3* mutants resembles the striated SC-CR staining in polycomplexes, relating to Figure 3**

(A) STED images of polycomplexes from *cohesin(-)* and *htp-3(-)* worms, stained for the SC-CR components SYP-2 or SYP-5, or the axis component HTP-3. In all cases, regularly spaced striations are observed. Graphs of line scans (yellow arrows) are shown below the images. (B) Quantification of the inter-peak distances in the line scans in Panel A shows spacing resembling an assembled synaptonemal complex. The inter-peak distance in wild-type animals from Figure 3H is shown for reference. (C) STED images of SC-CR aggregates in *him-3*$^{KK170\text{-}171EE}$ animals stained against SYP-2. While not showing regular lamellae, the anisotropic staining is distinct from the single-lamella SC-CR staining in wild-type animals.

### Figure S6

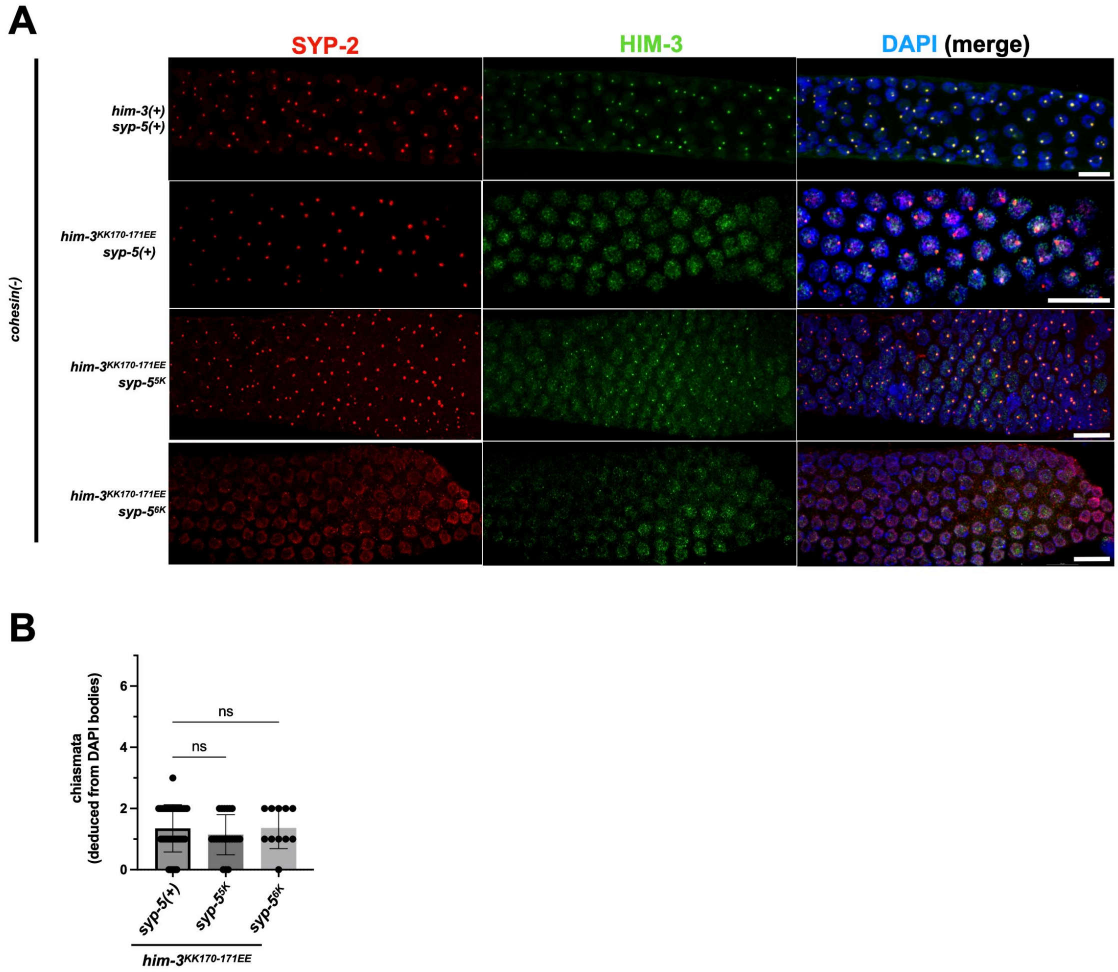


**Figure S6: Additional characterization of *him-3$^{KK170\text{-}171EE}$ syp-5$^{5K}$* and *him-3$^{KK170\text{-}171EE}$ syp-5$^{6K}$* worms, relating to Figure 4**

(A) The pachytene region of gonads from worms of the indicated genotypes, corresponding to Figure 4B, stained for the SC-CR component SYP-2 (red) and the axis component HIM-3 (green). The merged images on the right also show DNA (DAPI, blue). Scale bars = 10 μm. (B) Chiasmata number in worms of the indicated genotypes, deduced from the number of DAPI bodies at diakinesis.

### Figure S7

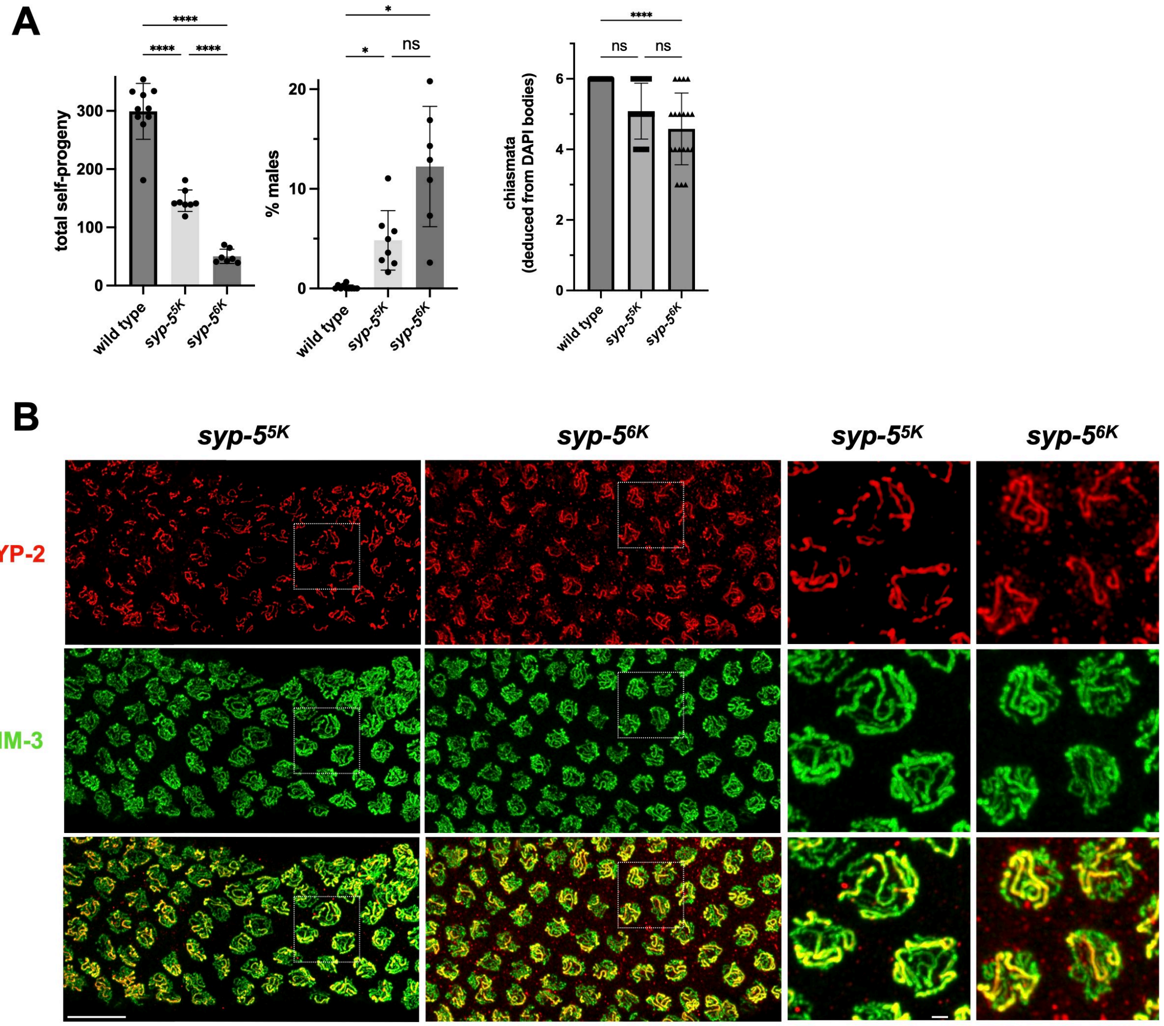


**Figure S7: Meiotic phenotypes *syp-5^5K^* and *syp-5^6K^* worms, relating to Figure 4**

(A) Left, total self-progeny from hermaphrodites of the indicated genotypes. Middle, percentage of males among self-progeny of hermaphrodites of the indicated genotypes, indicative of meiotic *X* chromosome non-disjunction. Right, chiasmata number deduced from the number of DAPI bodies at diakinesis. Wild-type values, taken from Figure 3, are shown for reference. (B) Pachytene nuclei stained for the SC-CR component SYP-2 (red) and the axis component HIM-3 (green), with merged images shown on the bottom. Note the extensive asynapsis in both mutants (i.e., axes lacking SC-CR staining). Scale bars = 10 µm for the gonad views 1 µm for the zoomed-in views on the right.

### Figure S8

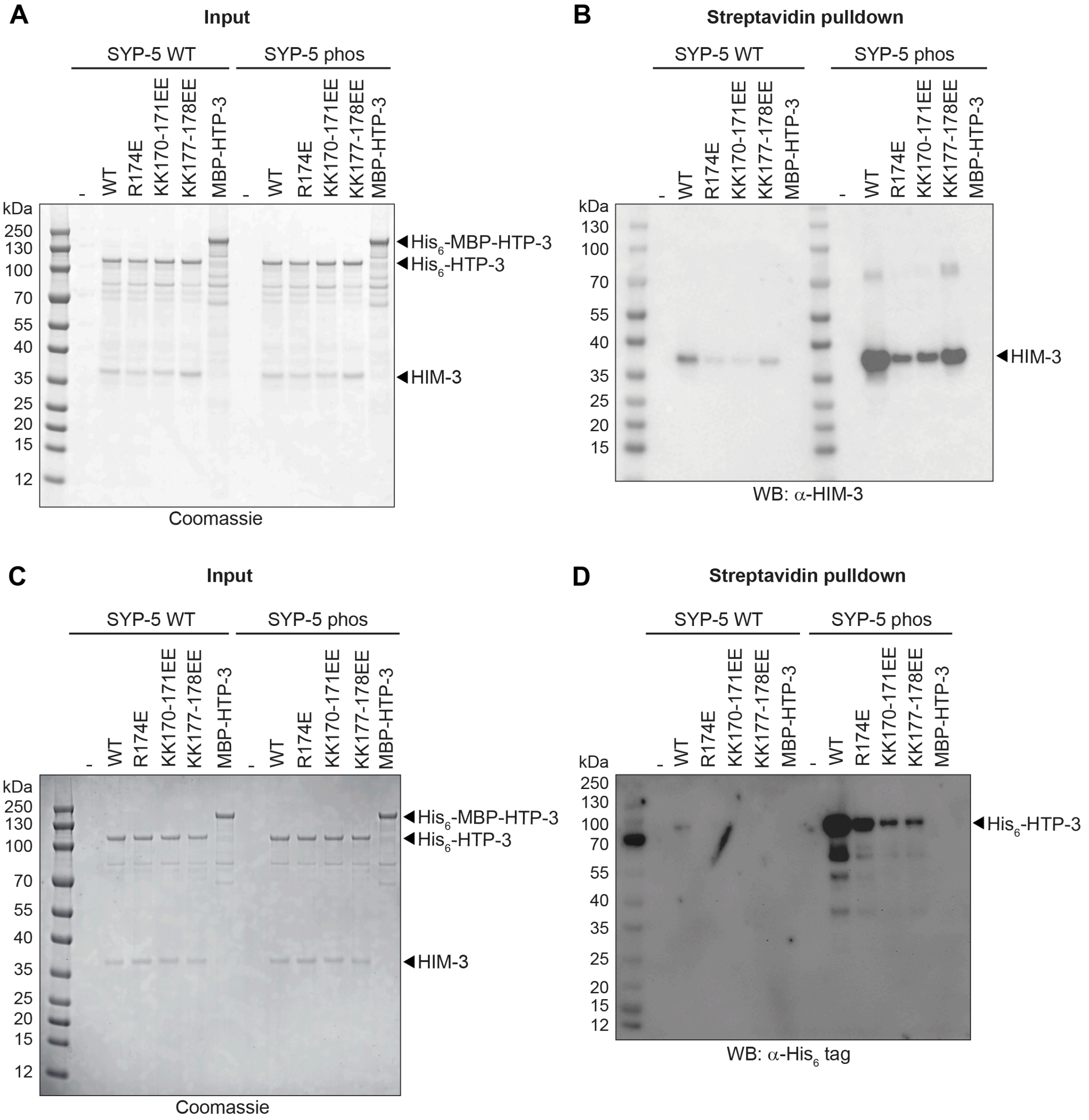


**Figure S8: Streptavidin pulldowns showing binding of HIM-3 to the SYP-5 C-terminus, relating to Figure 4J**

(A) Coomassie-stained SDS-PAGE gel showing input samples for streptavidin pulldowns with biotinylated SYP-5 peptides (SYP-5 WT: SYP-5 residues 528-547; SYP-5 phos: SYP-5 residues 528-547 with phosphorylated S541). Proteins used were $His_6$-HTP-3–HIM-3 complexes (wild type or the indicated mutants in the HIM-3 positive patch) or $His_6$-MBP-HTP-3 alone (MBP-HTP-3). (B) Anti-HIM-3 western blot of SDS-PAGE gel showing streptavidin-bound samples from the samples in panel (A). (C) As panel (A). (D) Anti-$His_6$-tag western blot western blot of SDS-PAGE gel showing streptavidin-bound samples from the samples in panel (C).

### Figure S9

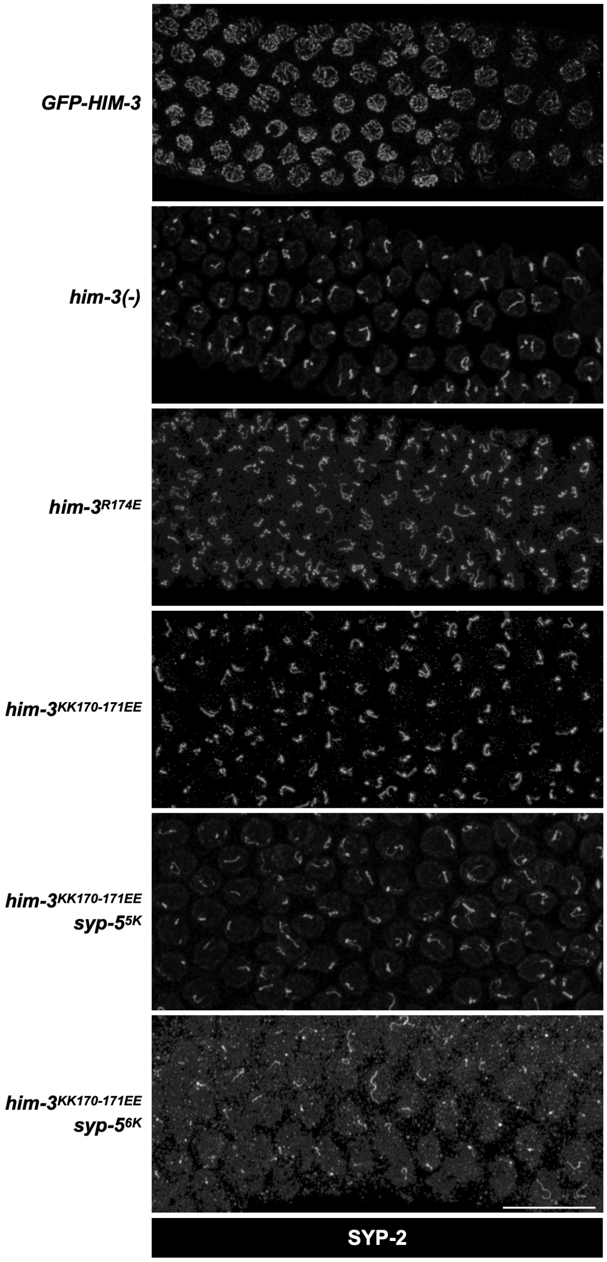


**Figure S9: SC-CR staining in various genotypes used to quantify the number of synapsed chromosomes, relating to Figures 3 and 4**

The pachytene region of gonads from worms of the indicated genotypes used to quantify the number of synapsed chromosomes (SC-CR threads per nucleus). Gonads were stained for the SC-CR component SYP-2. Scale bars = 10 μm.

### Figure S10

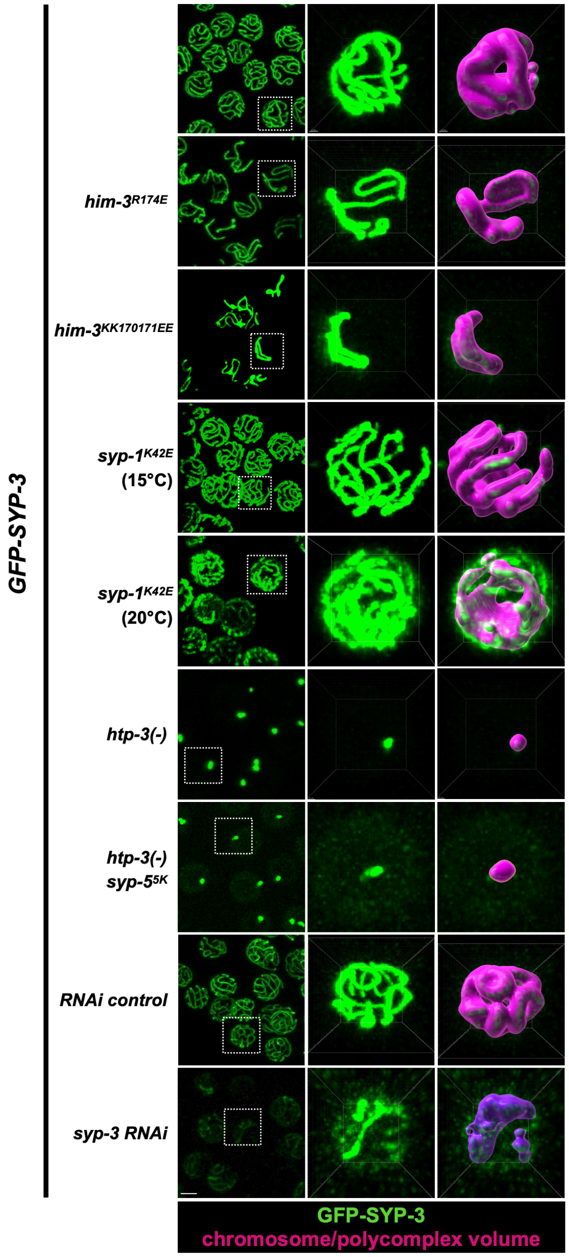


**Figure S10: GFP-SYP-3 abundance and localization, relating to Figures 5 and 6**
Pachytene nuclei from live worms expressing GFP-SYP-3. The middle column zooms in on a single nucleus. Right, Surface rendering of SC-CR structures, either synapsed chromosomes, or, in *htp-3(-)* animals, polycomplexes.

### Figure S11

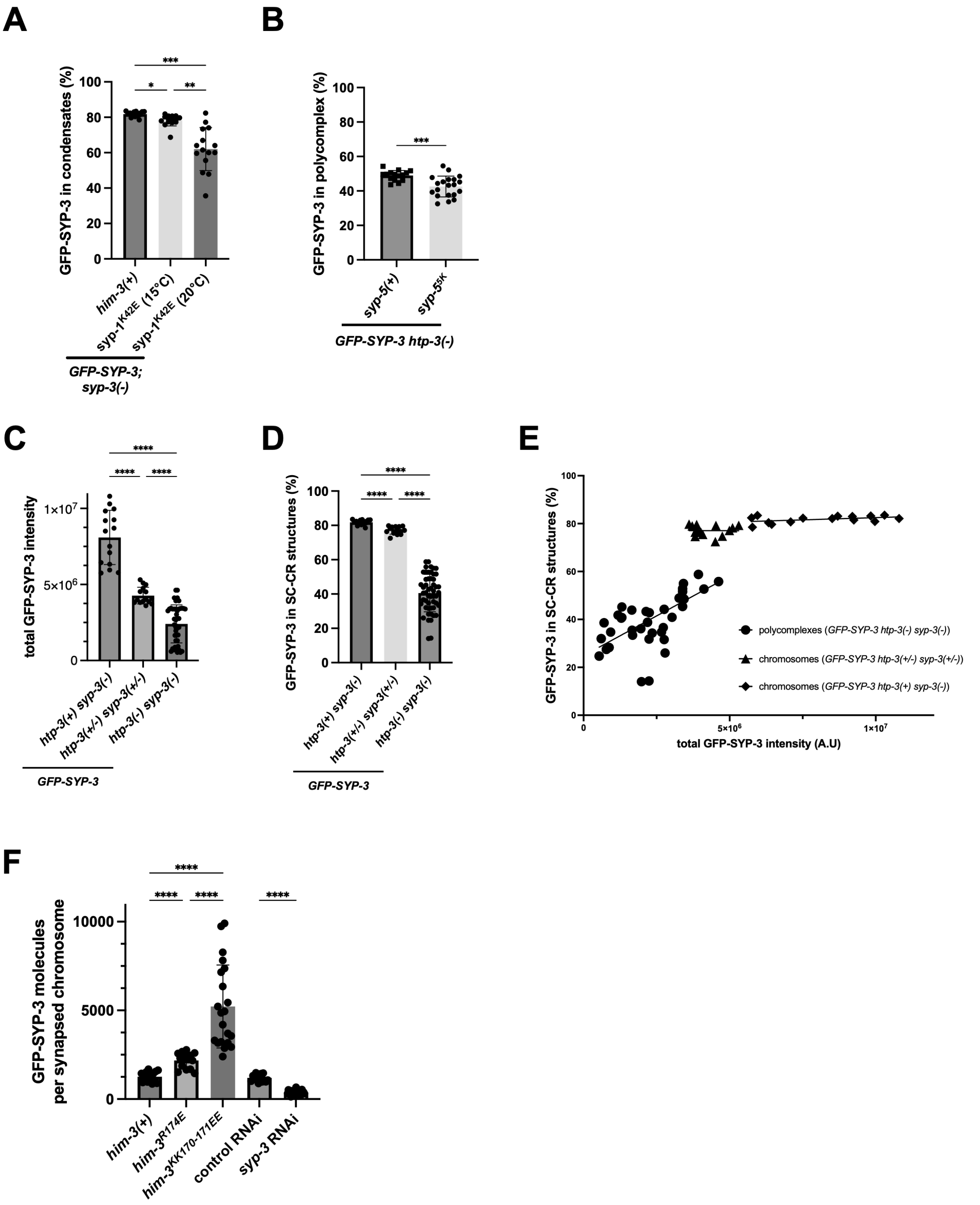
A
***
*
**
GFP-SYP-3 in condensates (%)
100
80
60
40
20
0
him-3(+)
syp-1K42E (15°C)
syp-1K42E (20°C)
GFP-SYP-3;
syp-3(-)
B
***
GFP-SYP-3 in polycomplex (%)
100
80
60
40
20
0
syp-5(+)
syp-55K
GFP-SYP-3 htp-3(-)
C
****
****
****
total GFP-SYP-3 intensity
1×10^7
5×10^6
0
htp-3(+) syp-3(-)
htp-3(+/-) syp-3(+/-)
htp-3(-) syp-3(-)
GFP-SYP-3
D
****
****
****
GFP-SYP-3 in SC-CR structures (%)
100
80
60
40
20
0
htp-3(+) syp-3(-)
htp-3(+/-) syp-3(+/-)
htp-3(-) syp-3(-)
GFP-SYP-3
E
GFP-SYP-3 in SC-CR structures (%)
100
80
60
40
20
0
0
5×10^6
1×10^7
total GFP-SYP-3 intensity (A.U)
polycomplexes (GFP-SYP-3 htp-3(-) syp-3(-))
chromosomes (GFP-SYP-3 htp-3(+/-) syp-3(+/-))
chromosomes (GFP-SYP-3 htp-3(+) syp-3(-))
F
****
****
****
****
GFP-SYP-3 molecules
per synapsed chromosome
10000
5000
0
him-3(+)
him-3R174E
him-3KK170-171EE
control RNAi
syp-3 RNAi

**Figure S11: Quantification of GFP-SYP-3 abundance and localization, relating to Figures 5 and 6**

(A) *syp-1*$^{K42E}$ is a temperature-sensitive mutation that exhibits increasing destabilization of the SC-CR with increasing temperature (Gordon *et al.* 2021). *syp-1*$^{K42E}$ worms exhibit very mild phenotypes at 15°C (permissive temperature), with significant synapsis defect at 20°C (semi-permissive temperature). The fraction of GFP-SYP-3 on chromosomes in animals of the indicated genotypes is significantly lower in *syp-1*$^{K42E}$ mutants. (B) The fraction of GFP-SYP-3 in polycomplexes (*htp-3(-)* worms) in *syp-5(+)* and *syp-5*$^{5K}$ background. (C) Quantification of the total amount of GFP-SYP-3. Note that GFP-SYP-3 is expressed in lower levels in the presence of the wild-type copy of *syp-3* (in the *htp-3(+/-) syp-3(+/-)* animals), as has been observed previously (Rog and Dernburg 2015). The *htp-3(-)* animals are progeny of trans-heterozygous animals (*htp-3(+/-) syp-3(+/-)*), likely accounting for the lower GFP-SYP-3 expression. (D) The percentage of GFP-SYP-3 in SC-CR structures (chromosomes or polycomplexes) is dramatically lower for polycomplexes versus chromosomes. (E) Enrichment of GFP-SYP-3 on SC-CR structures (polycomplexes or chromosomes) as a function of total GFP-SYP-3 levels. Circles, *GFP-SYP-3 htp-3(-) syp-3(-)* polycomplexes; triangles, *GFP-SYP-3 htp-3(+/-) syp-3(+/-)* chromosomes; diamonds, *GFP-SYP-3 htp-3(+) syp-3(-)* chromosomes. Note the lower enrichment of GFP-SYP-3 on polycomplexes versus chromosomes when comparing animals with similar expression levels. (F) The number of GFP-SYP-3 molecules per synapsed chromosome, calculated based on the average number of synapsed chromosomes and the chromosome-associated GFP-SYP-3 fluorescence. *him-3*$^{R174E}$ and *syp-3* RNAi animals exhibited ~two-fold increase and decrease, respectively, in the number of GFP-SYP-3 molecules per synapsed chromosomes. These numbers are in line with the ~two-fold variability during meiotic progression under physiological conditions (Pattabiraman *et al.* 2017). The much larger amount of GFP-SYP-3 per chromosome in *him-3*$^{KK170-171EE}$ animals is consistent with the much thicker SC-CR in this mutant (Figure 3F).

### Figure S12

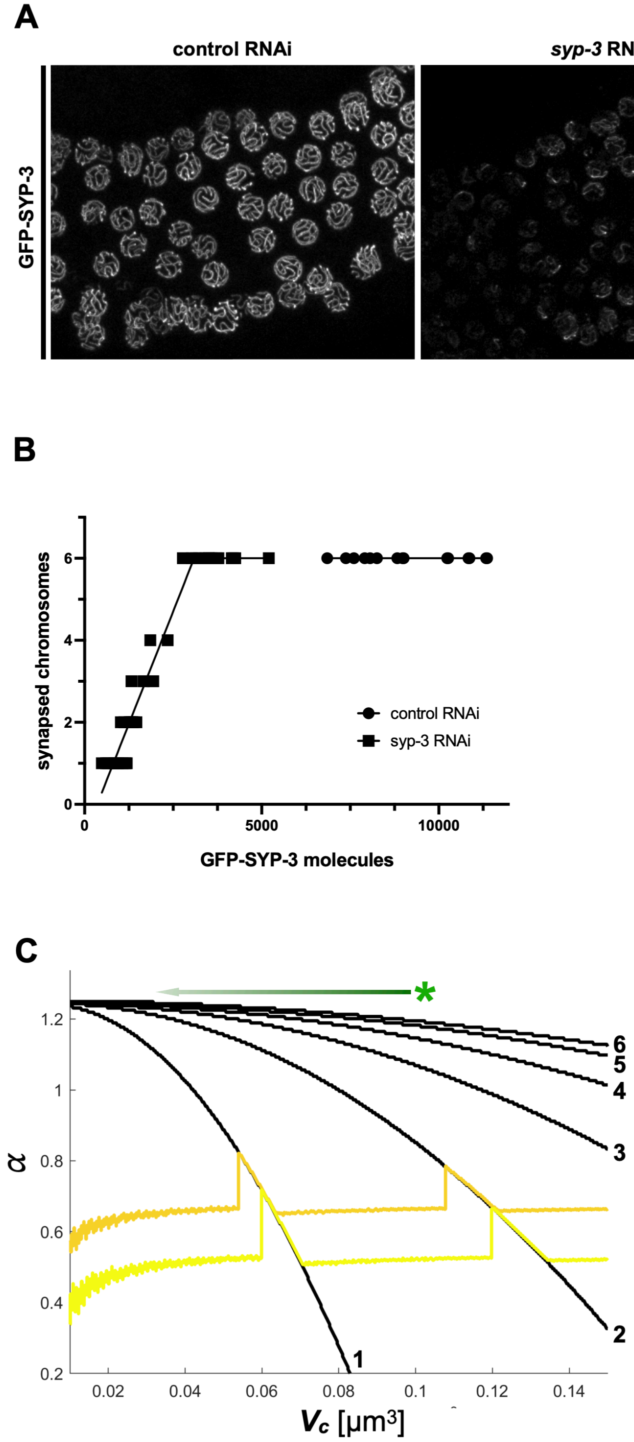
A
control RNAi
syp-3 RNAi
syp-3 RNAi (overexposed)
GFP-SYP-3
B
synapsed chromosomes
0
2
4
6
0
5000
10000
GFP-SYP-3 molecules
control RNAi
syp-3 RNAi
C
α
1.2
1
0.8
0.6
0.4
0.2
0.02
0.04
0.06
0.08
0.1
0.12
0.14
Vc [µm3]
1
2
3
4
5
6

**Figure S12: Meiotic outcomes of reduced SC-CR levels, relating to Figure 6**
(A) The pachytene region of the gonad in live worms expressing GFP-SYP-3. Worms were treated with RNAi control (empty plasmid) or RNAi against *syp-3* (Libuda *et al.* 2013). The left and middle images were taken with the same fluorescence and detector settings. The image on the right is overexposed to highlight the few chromosomes undergoing synapsis upon SC-CR depletion. Scale bar = 1 μm. (C) Quantification of the experimental data in panel B. The number of synapsed chromosomes upon lowering of SC-CR levels (circles, control RNAi; squares, *syp-3* RNAi). The number of GFP-SYP-3 molecules was determined by dividing the total nuclear fluorescence by the fluorescence of a single GFP-SYP-3 molecule (calculated based on 9,200 molecules in the control RNAi condition). Each point indicates a single nucleus. (C) The contour plot from Figure 6B is overlayed with the effect of reducing the number of SC-CR molecules (green arrow). The wild-type scenario is shown with a green asterisk. See text for details.

### Figure S13

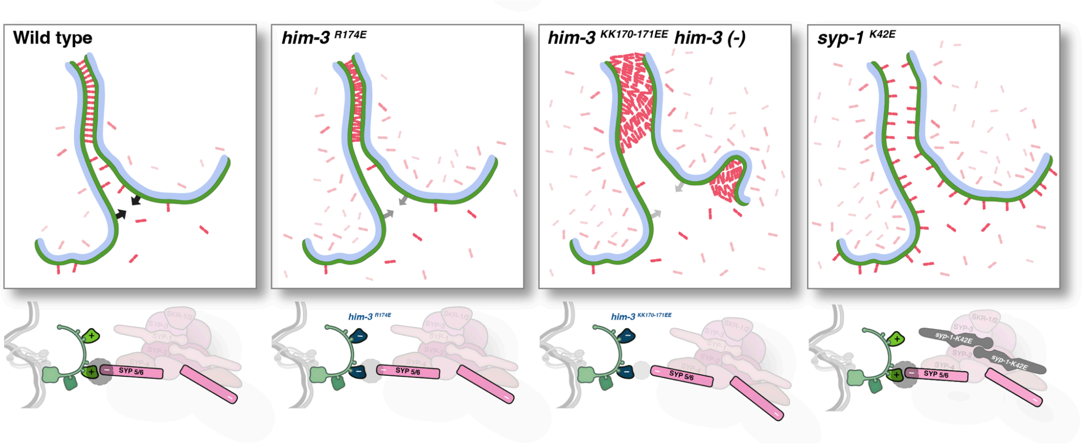


**Figure S13: Graphical representation of the mutant phenotypes used in this study, relating to Figure 6**

Depiction of mutant scenarios, with SC-CR subunits shown in red/magenta, axes shown in green, and the chromosomes represented in light blue. See text for details.
