## Supplementary Note 1 for "The synaptonemal complex aligns meiotic chromosomes by wetting"

### SUPPLEMENTARY NOTE 1: BIOPHYSICAL MODELING OF SYNAPTONEMAL COMPLEX ASSEMBLY

#### 1. QUANTITATIVE CONSIDERATIONS OF SYNAPTONEMAL COMPLEX STRUCTURE

##### 1.1 Structural parameters of meiotic nuclei and the synaptonemal complex in *C. elegans*

Meiotic nuclei in the oogenic *C. elegans* gonad are ~3.5 μm in diameter (22.5 $\mu m^3$; see Fig. 5A for a table of key physical parameters). The center of each nucleus is occupied by a nucleolus, of ~2 μm diameter (Goldstein and Slaton 1982). We also assume that chromatin occupies about 50% of the nucleus. This leaves a nucleoplasmic volume of 9.2 $\mu m^3$.

We estimated the number of HIM-3 molecules based on the binding of an average of eight HIM-3 molecules to the base of the ~160kb chromatin loops (Woglar et al. 2020), for a total of 11,000 molecules per nucleus (using a 100Mb haploid genome size). SC-CR components are notoriously challenging to functionally tag with fluorescent proteins. Given that all SC-CR components in worms are co-dependent for their expression (Gordon and Rog 2023; MacQueen et al. 2002; Colaiácovo et al. 2003; Smolikov, Schild-Prüfert, and Colaiácovo 2009; Smolikov et al. 2007; Hurlock et al. 2020; Zhang et al. 2020), we estimated the number of molecules of the SC-CR component SYP-3, for which a functional tagged version exists (Rog and Dernburg 2015). We compared, under identical imaging conditions, the nuclear fluorescence of worms expressing either GFP-HIM-3 or GFP-SYP-3, revealing there are 9,200 SYP-3 molecules per nucleus (Fig. 5C). To connect to our SYP-5-based model, we assume 1:1 stoichiometry between SYP-5 and SYP-3 protein, consistent with recent *in vitro* data showing four each of SYP-3 and SYP-5 in a recombinantly expressed SC-CR (Blundon et al. 2024).

Importantly, these measurements, as well as all other quantitative measurements used for the model, rely on live imaging of fluorescently tagged components. This strategy avoids the challenges and assumptions associated with fixation and the use of antibodies.

##### 1.2 Phase separation of SC-CR components

When SC-CR components are not adsorbed onto the axes (e.g., in *htp-3* worms), they form nucleoplasmic assemblies called polycomplexes, which resemble in appearance stacked SC-CR ladders (Fig. S5 and (Hughes and Hawley 2020; Rog, Köhler, and Dernburg 2017)). Polycomplexes exhibit liquid-like properties, including spherical morphology and resorption upon merging (Rog, Köhler, and Dernburg 2017). These observations provide evidence that SC-CR components have an inherent tendency to phase separate in the nucleoplasm. Polycomplexes have been observed in many organisms, suggesting this is a conserved property of SC-CR components (Hughes and Hawley 2020).

We can use the volume of the polycomplex (~0.05 $\mu m^3$ ~ 0.54% of the nucleoplasm, Fig. 5E) and the fraction of SYP-3 in polycomplexes (~40%, Fig. S10B) to estimate that the enrichment (or partition coefficient) of SC-CR components in polycomplexes is 123 (40%/0.54%) / ([100% - 40%]/[100%-0.54%])). This number is similar to what is calculated based on immunofluorescent images (Figs. 1 and 2). We note that this value is higher than the *in vivo* partition coefficient measured for other condensates in *C. elegans*, such as P granules (partition coefficient for PGL-1: 12-36; (Fritsch et al. 2021)) or the centrosome (17 and 23 for SPD-2 and ZYG-9, respectively; (Woodruff et al. 2017)), suggesting that the self-interactions among SC-CR components may be stronger compared to those in the aforementioned biomolecular condensates.

Further, we can estimate the SYP-3 concentration in the condensate, $c_{cond}$, using the following equation that relates the number of SYP-3 molecules in the dilute and condensed phase, to the total number of SYP-3 molecules in the nucleoplasm, which is 9200 (Fig. 5C):

$$0.05\mu m^3 \times c_{cond} + (9.2 - 0.05)\mu m^3 \times \frac{c_{cond}}{123} = 9200. \qquad \text{[Eq. (1)]}$$

From Eq. (1), we find

$$c_{cond} \approx 74500 \mu\text{m}^{-3}. \quad \text{[Eq. (2)]}$$

In the analysis below, we denote the attractive binding energy between SC-CR components by $e_{SS}$. Notably, this binding energy combines multiple molecular events that promote interactions between SC-CR components. These interactions collectively result in the formation of ladder-like SC-CR structures, both between axes and in polycomplexes.

**1.3 The axes contain a fixed number of SC-CR binding sites**

Axes assemble onto chromosomes prior to, and independently of, the SC-CR (Zickler and Kleckner 2023; Gordon and Rog 2023). Axes then serve as a platform for the assembly of the SC-CR, indicating the inherent affinity between components of these two elements of the synaptonemal complex. We define the binding energy between the axis and the SC-CR as $e_{SH}$. As with $e_{SS}$, $e_{SH}$ denotes the ensemble of molecular interactions between axes and SC-CR components.

In Figs. 1-4, we provide evidence that an interface between a positive patch on the axis component HIM-3 and the negative C-terminus of the SC-CR component SYP-5 constitutes one of these molecular interactions. By mutating these domains, we weaken this interface, and by combining charge-swap mutations we were able to partially restore it. Our data also point to the existence of additional interfaces that mediate axis-SC-CR interactions. Below we use quantitative measurements in worms carrying the mutations we generated to develop and test a biophysical model of synaptonemal complex assembly.

Our measurements on the effects of mutations in *him-3* have an additional important implication. We observe a similar number of HIM-3 molecules on chromosomes even when we dramatically weaken axis-SC-CR interactions (Fig. S4B-D). Given that the SC-CR has an affinity to the axes, this is a non-trivial observation. A naïve model would predict that an assembled SC-CR would recruit more axes components, resulting in fewer diffuse HIM-3 molecules.

Our interpretation of this result is that axes components interact with chromosomes with a binding energy that is much stronger than $e_{SH}$. Even with such strong interaction, only 70% of HIM-3 molecules are on chromosomes (Fig. S4D). This suggests that the number of HIM-3 molecules on the chromosomes is limited due to structural considerations, and, in turn, that there is also a limited number of SC-CR binding sites on the axes. This latter idea is incorporated into the model.

These ideas are also hinted at by previously published data. For instance, conditions that prevent pairing and synapsis on a subset of chromosomes in worms (e.g., mutations in *him-8* or *zim-1-3*) do not cause a depletion of axes components on the asynapsed chromosome (Phillips et al. 2005; Phillips and Dernburg 2006). Similar observations have been made in mice (e.g., (Baudat et al. 2000)). In addition, the polycomplexes that form in budding yeast when chromosome pairing is prevented do not deplete axis components away from the chromosomes (Lin et al. 2010). Finally, even a relatively minor reduction in the number of HIM-3 molecules on chromosomes (caused by mutating one or two of the four HIM-3-binding sites on the component that recruits it to chromosomes, HTP-3) results in drastic effects on the kinetics and extent of synapsis (Kim, Kostow, and Dernburg 2015).

**1.4 Volume of SC-CR condensates**

Condensation is caused by attractive interactions among the SC-CR components (i.e., $e_{SS}$), and condensate volume is expected to increase as the strength of the binding energy, $e_{SS}$, increases (given constant number of SC-CR molecules). However, the total condensate volume is also affected by the condensate's interaction with its environment, such as wetting onto existing structures. Generally, wetting occurs due to the reduction of free energy via adhesive interaction (adsorption). As a result of this reduction of free energy, condensate volume is also expected to increase.

This is what we observe experimentally by measuring the fraction of SC-CR material in condensates, which serves as a proxy for volume (Fig. S11C-E). Specifically, we observe ~40% of SC-CR material in condensates that form purely through condensation (i.e., polycomplexes), versus ~80% of SC-CR material in condensates that form by relying on adhesion to axes *and* condensation (i.e., the wild-type scenario). Furthermore, in conditions that reduce either intra-SC-CR affinity (weaker $e_{SS}$; *syp-1*$^{K42E}$ or *syp-5*$^{5K}$) or axis-SC-CR affinity (weaker $e_{SH}$; *him-3*$^{R174E}$ or *him-3*$^{KK170\text{-}171EE}$), condensate volume is lower (Fig. S11A-B).

#### 2. SYNAPTONEMAL COMPLEX ASSEMBLY IN UNPERTURBED CONDITIONS

##### 2.1 Developing a free energy model

Given SC-CR components' tendency to phase separate and to interact with the axes, we postulate that SC-CR assembly onto the axes (synapsis) is driven by two processes: 1) binding to HIM-3 (with binding energy $e_{SH}$), and 2) minimization of interfacial energy in SC-CR condensates. Synapsis likely involves binding of some SC-CR components to HIM-3 on the axes, mediated by relatively strong binding energy $e_{SH}$, and condensation of the rest of SC-CR components, from the soluble pool and potentially from polycomplexes, onto the existing synapsed chromosomes to minimize the interfacial energy (due to surface tension).

These ideas are consistent with many previous observations, including: 1) upon over-expression of SC-CR in yeast, SC-CR components are continuously recruited onto already-synapsed chromosomes (Voelkel-Meiman et al. 2012). 2) Polycomplexes do not co-exist with synapsed chromosomes under most conditions; rather, they assemble prior to synapsis, following synaptonemal complex disassembly, or when SC-CR components are over-expressed (Hughes and Hawley 2020; Rog, Köhler, and Dernburg 2017; Goldstein and Slaton 1982). 3) SC-CR assembly is cooperative, meaning that under limiting SC-CR component numbers, a subset of chromosomes is completely synapsed rather than all chromosomes being partially synapsed (Hayashi, Mlynarczyk-Evans, and Villeneuve 2010).

We will now use a free energy model to account for the salient experimental observations. Our use of a free energy model is motivated by the formation of droplet-like polycomplexes by phase separation, which is a thermodynamic process that is accounted for by the minimization of free energy (Barrat and Hansen 2011). While the material state of chromosome-associated SC-CR is not as well defined, SC-CR components exchange dynamically between synapsed chromosomes and the nucleoplasm, as indicated by photoactivation and photobleaching experiments (Pattabiraman et al. 2017; Nadarajan et al. 2017; Rog, Köhler, and Dernburg 2017), and they diffuse laterally within the SC-CR (von Diezmann, Bristow, and Rog 2024). Furthermore, SC-CR components, in worms and in other organisms, do not harbor a nucleotide hydrolysis domain, which suggests that assembly is not an out-of-equilibrium process, and, unlike polymerization of actin filaments and microtubules, it does not require NTP hydrolysis. Together, these observations suggest that molecular interactions underlying synaptonemal complex assembly are reversible at relevant timescales. Hence, local binding and unbinding events, when averaged among many molecules, could allow sampling of the local energy landscape, motivating again the use of our free-energy-based model.

Our simple free energy model is as follows:

$$F_{tot} = e_{SH} n_{SH}^{bond} N_{syn-chr} + F_A \qquad \text{[Eq. (3)]}$$

where $n_{SH}^{bond}$ is the number of HIM-3--SYP-5 binding sites per pair of chromosomes (which we can estimate to be $\sim 500$ ($2 \times \frac{6000nm}{24.2nm} = 496$, 2 for the bilayer [i.e., head-to-head] structure, 6000nm being the length of a synapsed chromosome [the six worm chromosomes are similar in size], and 24.2nm being the spacing between the ladder rungs observed in EM images of the SC-CR (Fig. 5)); notably, this number is very similar to the independently-derived minimal number of SYP-3 molecules required to synapse chromosomes, as measured by RNAi experiments, below), $N_{syn-chr}$ is the number of synapsed chromosomes, and $F_A$ is the interfacial free energy due to the presence of the SC-CR-nucleoplasm interface, which we model as

$$F_A = A\left[1 + \frac{b(w-w_0)^2}{w_0^2}\right]\varsigma e_{SS} \qquad \text{[Eq. (4)]}$$

where $\varsigma e_{SS}$ is the surface tension written in this form because it is expected to be proportional to $e_{SS}$ - the self-interaction energy between SC-CR components, $A\left[1 + \frac{b(w-w_0)^2}{w_0^2}\right]$ models the interfacial area per synapsed chromosome that depends on the thickness $w$ of the adsorbed condensate. Note that we have assumed that the surface area depends on the thickness of the adsorbed condensate in such a way that there is an optimal thickness $w_0$, with $A$ being the surface area: $N_{syn-chr} \times (6000nm \times 2 \times 100nm) + [N_{syn-chr}] \times 2 \times w \times 100nm$ (i.e., the areas of 4 sides of a rectangular prism), when $w = w_0$, and $b$ being a phenomenological parameter. (See Section 4, below, for robustness analysis of these and other parameters.)

Such a dependency can be rationalized, for example, by appreciating that the interfacial profile can adopt a concave or convex profile (in a head-on view) depending on the interfacial structure at the molecular scale (Model Fig. 1). However, we note that as the adsorption corresponds to a bilayer structure of SC-CR components, our use of interfacial energy should be viewed as an approximation that aims to account for the loss of self-interactions at the periphery of the condensate.

---

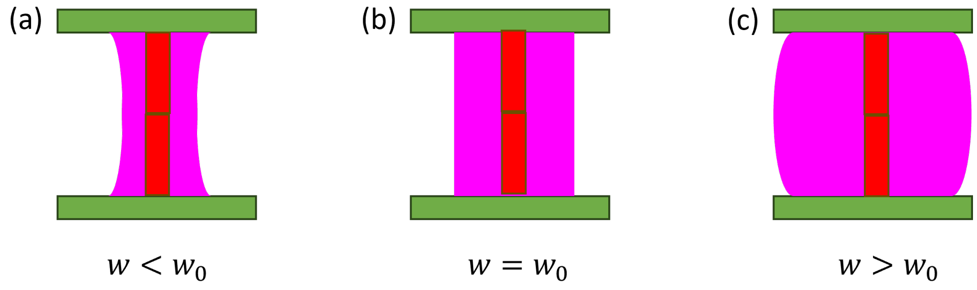


**Model Figure 1: Illustrative diagrams of cross sections of synapsed chromosomes.** Axes are shown as green bars, with different amounts of SC-CR molecules per condensate (magenta). The central pillars (red) depict the ladder-like structure of the SC-CR, as well as the number of SC-CR binding sites on the axes. The exposed surface area of the SC-CR is larger when the thickness of the SC-CR, $w$, is either smaller (a; concave morphology) or larger (c; convex morphology) than the optimal value ($w_0$, shown in b).

---

We can estimate the overall condensate volume, $V_c$, by the following:

$$V_c = N_{syn-chr} \times w \times 6\mu m \times 100nm \qquad \text{[Eq. (5)]}$$

where $6\mu m$ is the length of a synapsed chromosome and 100nm is the distance between the two axes in synapsed chromosomes. Under wild-type conditions, $w$ is estimated to be 30-40nm (based on the distribution of SC-CR components in head-on view by super-resolution localization microscopy; (Köhler et al. 2017, 2020)). This results in a value of

$$V_c \approx 0.1\mu m^3 \qquad \text{[Eq. (6)]}$$

We note that this value is comparable, albeit slightly larger than the polycomplex volume in *htp-3* mutants (Section 1.1), as would be expected due to wetting of the axes (Section 1.3).

**2.2. Expected number of synapsed chromosomes**

We can minimize the total free energy $F_{tot}$ to reveal $N_{syn-chr}$ - the number of synapsed chromosomes. Since the SC-CR binds cooperatively to chromosomes (Hayashi, Mlynarczyk-Evans, and Villeneuve 2010), such occupied sites will be distributed on the smallest possible number of chromosomes. We can rewrite the total free energy as

$$F_{tot} = e_{SS}\left\{\alpha n_{SH}^{bond} N_{syn-chr} - N_{syn-chr} A\left[1 + \frac{b(w-w_0)^2}{w_0^2}\right]\varsigma\right\}, \quad \text{[Eq. (7)]}$$

where

$$\alpha = \frac{e_{SH}}{e_{SS}} \quad \text{[Eq. (8)]}$$

denotes the ratio between the axis-SC-CR binding energy and SC-CR self-interaction energy (both are negative because the interactions are attractive). Further, for a given condensate volume $V_c$, we can infer the thickness $w$.

For instance, if we fixed the condensate volume $V_c$ to be 0.1μm$^3$, we can minimize the free energy, $F_{tot}$, as a function of $\alpha = \frac{e_{SH}}{e_{SS}}$ to find the expected number of synapsed chromosomes, $N_{syn-chr}$ (Model Fig. 2). This analysis shows that as $\alpha$ is diminished, fewer chromosome pairs will synapse.

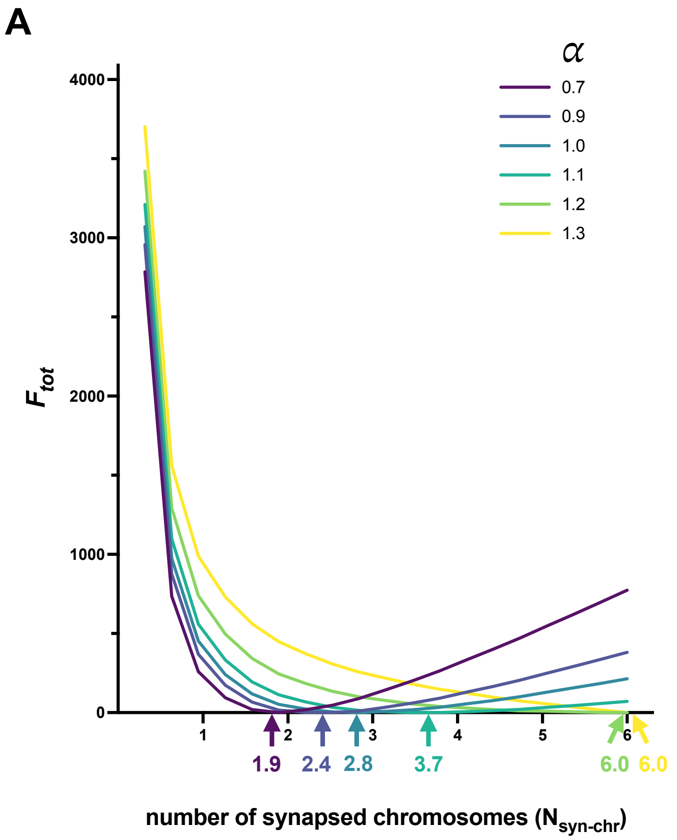


**Model Figure 2: Minimizing free energy $F_{tot}$ to predict the number of synapsed chromosomes $N_{syn-chr}$ with changing values of $\alpha = e_{SH}/e_{SS}$.** (A) Each curve represents the free energy $F_{tot}$ (in arbitrary units) for different value of $\alpha = e_{SH}/e_{SS}$. For each value, the curve's minimum reflects the predicted number of synapsed chromosomes $N_{syn-chr}$ (shown as a colored number below the x-axis). We see that as $\alpha = e_{SH}/e_{SS}$ changes from 0.7 to 1.4, the number of synapsed chromosomes increases from 1.9 to 6. (B) Summary of the results from (A), shown as a continuous curve linking different values of $\alpha = e_{SH}/e_{SS}$ to the number of synapsed chromosomes $N_{syn-chr}$. The dashed lines, corresponding to the wild-type ($N_{syn-chr} = 6$) and *him-3*$^{R174E}$ ($N_{syn-chr} = 3$) conditions, reveal $\alpha > 1.2$ and $\alpha = 1.0$, respectively. Model parameter: the number of HIM-3-SYP-5 binding sites, $n_{SH}^{bond}$, is 496 ($\sim 2 \times 6\mu m/22.4nm$), $A \sim 2 \times 100nm \times (6\mu m + w)$ (corresponding to the four lateral sides of the rectangular prism), $w_0 = 60nm$ (i.e., about twice the expected thickness in the wide type (Eq. (5))), $\zeta = 1/(100nm \times 24.2nm)$, which roughly estimates the inverse of the surface area per SYP-5 protein at the interface, and $b = 0.25$. [Data in Data_Model_Figure_2A.xlsx and Data_Model_Figure_2B.xlsx.]

---

More generally, we can investigate the expected number of synapsed chromosomes, $N_{syn-chr}$, as both $\alpha$ and $V_c$ vary. The results are shown in Model Fig. 3:

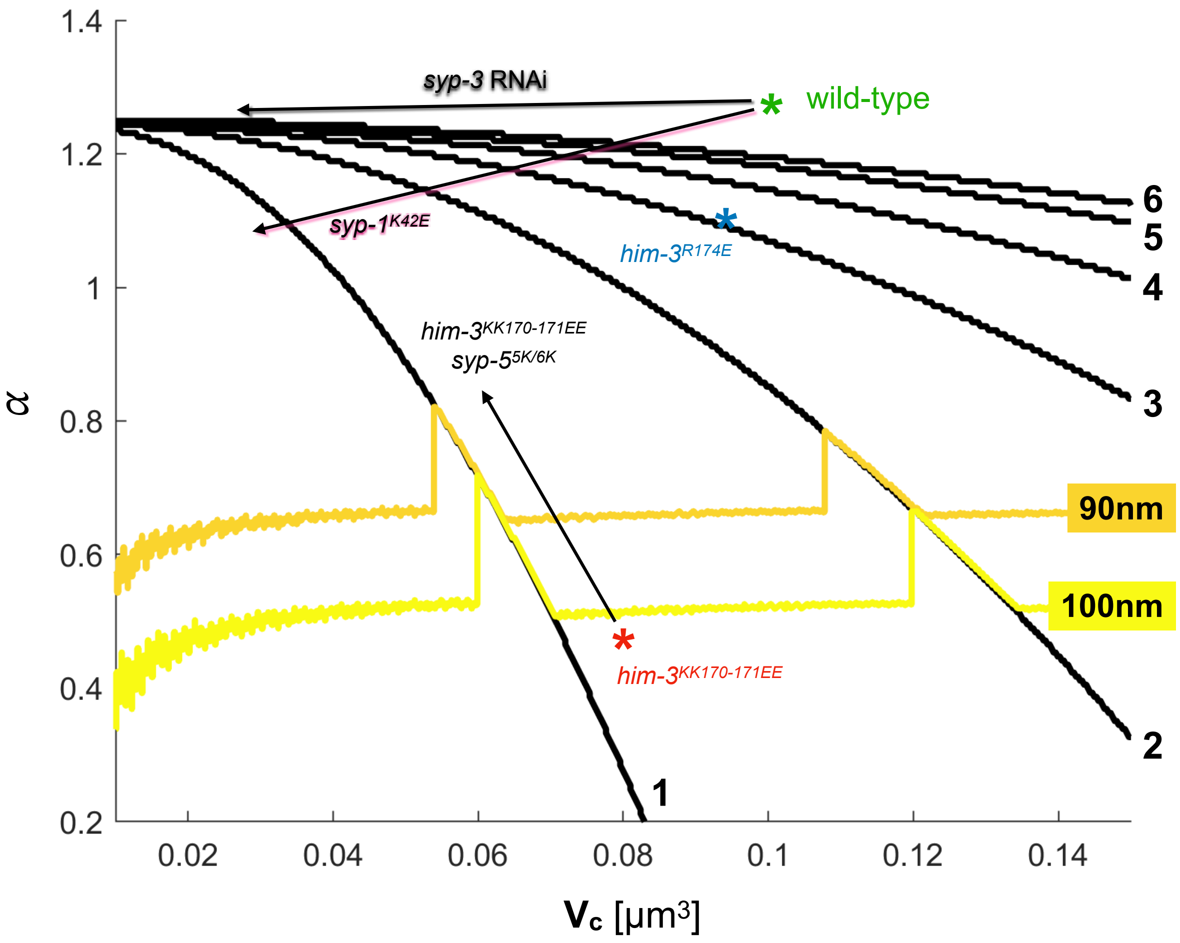


**Model Figure 3: Contour plot of the number of synapsed chromosomes as $\alpha$ and $V_c$ vary.** The black curves indicate the minimum values of $\alpha$ and $V_c$ necessary to synapse the indicated number of chromosomes. The yellow and orange lines denote the estimated thickness $w$ of the SC-CR (100nm and 90nm, respectively). Thickness larger than ~100nm is likely to be too big for a unilamellar SC-CR and thus a multi-lamellar SC-CR structure would be expected below the yellow-orange lines (referred to in Fig. 3 as "wide synapsis"). Green, blue and red asterisks denote wild-type, *him-3*$^{R174E}$ and *him-3*$^{KK170\text{-}171EE}$ worms, respectively. Black arrow depicts the transition from *him-3*$^{KK170\text{-}171EE}$ worms to *him-3*$^{KK170\text{-}171EE}$ *syp-5*$^{5K}$ and *him-3*$^{KK170\text{-}171EE}$ *syp-5*$^{6K}$ worms. The model parameters are the same as in Model Fig. 2. [Data in Data_Model_Figure_3.xlsx.]

The wild-type condition corresponds to $V_c \approx 0.1\mu m^3$ with all six chromosomes synapsed, which indicates that $\alpha >$ 1.2. Note that this value of $\alpha$ is a lower limit, as higher $\alpha$ values will also yield 6 synapsed chromosomes. As indicated by our *syp-3* RNAi experiment discussed below, $\alpha$ must be somewhat higher ($\alpha > 1.25$). The wild-type condition is indicated by a green asterisk in Model Fig. 3.

#### 3. MODEL PREDICTIONS IN PERTURBED CONDITIONS

##### 3.1 Impact of *him-3* mutations

*him-3*$^{R174E}$: This mutation results in ~3 synapsed chromosomes while the total condensate volume is only slightly less than the WT condition (92% of the wild-type; Fig. 5D; this is likely due to a weaker $e_{SH}$, which reduces adhesion energy of the condensate onto the axes). This translates to $\alpha = 1.0$. Those chromosomes that synapse will have a higher number of SC-CR components (since the total condensate volume, $V_c$, is barely changed), consistent with experimental observations (Fig. S11F). This condition is schematically indicated by the blue asterisk in Model Fig. 3.

*him-3*$^{KK170\text{-}171EE}$: Worms carrying this mutation exhibit 1-2 SC-CR bodies associated with axes, which we treat here as synapsed chromosomes, while $V_c$ is reduced to 81% of wild-type levels (Fig. 5D). This condition is schematically indicated by the red asterisk in Model Fig. 3.

We note that the inter-axes distance in this condition becomes variable and the SC-CR can exhibit multi-lamellar structures in certain segments (Fig. 3). This suggests that the amount of SC-CR material sandwiched between the axes is so large to the extent that the axes can no longer accommodate it. Hence, the condensate is forced to form a multi-lamellar structure. Specifically, we can estimate the maximum thickness of a single-lamellar SC-CR allowed on synapsed chromosomes as follows: the thickness of a single-lamellar SC-CR with 1.5 synapsed chromosomes for a condensate volume of ~0.08μm$^3$ (Eq. (8)) would be around 100nm ($\sim 0.08\mu m^3/(1.5 \times 6\mu m \times 100nm)$). The fact that we observe multi-lamellar structures suggests that the maximum possible thickness for a unilamellar SC-CR is somewhat lower than 100nm, but likely larger than 60-80nm. The latter number is derived from the roughly x2 accumulation of SC-CR components on chromosomes throughout meiotic prophase in wild-type meiosis, where aberrant SC-CR structures are not observed (Pattabiraman et al. 2017).

##### 3.2 Impact of perturbing SC-CR self-interactions

We examine two conditions that perturb SC-CR self-interaction (i.e., $e_{SS}$). The first is mutating *syp-5* (*5K* and *6K* mutations) in the *him-3*$^{KK170\text{-}171EE}$ background. These mutations do not increase the number of synapsed chromosomes (still, only about 1-2), but they help restore SC-CR morphology closer to normal thickness (unilamellar SC-CR). Our analysis using polycomplexes suggests that these mutations restore some of the affinity between SYP-5 and HIM-3 (Fig. 4), as well as reduce the self-interaction between SC-CR components resulting in smaller condensates (for *syp-5*$^{5K}$, 25% reduction in $V_c$; Fig. S11B) or a failure to form condensates (for *syp-5*$^{6K}$; Fig. 4). In the context of the model, $\alpha$ in the double-mutants is larger than in the single *him-3* mutant (larger $e_{SH}$ and smaller $e_{SS}$) and $V_c$ is smaller (fewer molecules in condensates). Plotting these changes as a vector on Model Fig. 3 yields a diagonal upward-left vector (black arrow). Importantly, this vector brings the thickness of the condensate below the threshold of multi-lamellar synapsis (yellow-orange lines), consistent with our empirical observations.

The second condition is the *syp-1*$^{K42E}$ mutation. This is a temperature-sensitive mutation in a region of SYP-1 located in the middle of the SC-CR (Gordon et al. 2021). This mutation affects $V_c$ (Fig. S11). This is expected given the smaller and less abundant polycomplexes that form at 20°C (semi-permissive temperature) and the lack of polycomplexes at 25°C (where worms are essentially sterile; Fig. 4 in (Gordon et al. 2021)). This mutation also affects the number of synapsed chromosomes, with close to 6 chromosomes at 15°C, ~4 synapsed chromosomes at 20°C, and only about 0-1 synapsed chromosomes at 25°C (Figs. 1-2 in (Gordon et al. 2021)). Based on the location of the mutation, the effects on polycomplex formation and hypersensitivity to hexanediol (Fig. 4 in (Gordon et al. 2021)), this mutation is likely to affect $e_{SS}$. However, it is possible that it also affects $e_{SH}$ indirectly, by impacting the extent of cooperativity between SC-CR subunits. Such an effect is consistent with the loss of the 'bilayer' (head-to-head) morphology of the SC-CR (Almanzar et al. 2023). Overlayed on the $\alpha$-to-$V_c$ diagram, the effects of *syp-1*$^{K42E}$ with increasing temperature resemble a left-pointing, mostly horizontal vector with increasing temperature (black/pink arrow in Model Fig. 3).

##### 3.3 Impact of reducing the availability of SC-CR components

To gauge the strength of HIM-3-SYP-5 binding, we use *syp-3* RNAi, which leads to a variable reduction in the number of SC-CR subunits in the nucleus (Fig. S12; (Hayashi, Mlynarczyk-Evans, and Villeneuve 2010; Libuda et al. 2013)). We measured, in parallel, the number of SYP-3 molecules and the number of synapsed chromosomes. Since the molecular components are not altered, such a reduction would mostly affect $V_c$. These experiments indicated that all six pairs of chromosomes synapse even when the number of SC-CR molecules is reduced to around 2500 (Figure 6C; 70-80% reduction relative to wild-type levels; (Libuda et al. 2013)). On Model Fig. 3, the *syp-3* RNAi experiment reflects a mostly horizontal line, with constant $\alpha$ and decreasing $V_c$ (shown as a black line). The ability to synapse all chromosomes at these very low $V_c$ values ($V_c \approx 0.02 - 0.03\mu\text{m}^3$) indicates that $\alpha >$ 1.25 (rather than $\alpha > 1.2$, as was deduced based solely on analysis of the wild-type scenario; Model Fig. 2). Importantly, the empirically derived minimal number of SC-CR molecules necessary to synapse all chromosomes (~2500) lends support to our previously calculated number of available binding sites on the axes of the six chromosomes ($6 \times \sim 500 = \sim 3000$).

Under such an extreme reduction in SC-CR numbers, there is likely to be only a single 'sheet' of SC-CR molecules, and accordingly, the use of the term condensate is no longer appropriate. In this condition, SC-CR-axis association is better described as an adsorption process at the molecular scale. Below 2,500 SC-CR molecules, the number of synapsed chromosomes decreases linearly with a further decrease in SC-CR molecule number (Fig. S12B), consistent with an adsorptive process that is limited by the number of SC-CR molecules.

#### 4. ROBUSTNESS ANALYSIS

We wanted to test the robustness of our model to measurement errors and to a variety of parameters. Several of the parameters and measurements used in our model would cause a minor effect on the exact value of $V_c$. These include the number of SC-CR molecules, the size of polycomplexes, the reduction in SC-CR numbers in the *syp-3* RNAi experiments, the dimensions of the SC-CR (ladder rung spacing and thickness) and the fraction of SC-CR molecules in condensates. Errors in these parameters are likely to cause a minor shift in the value of $V_c$, which translates to uncertainty in the x-axis location of the various experimental conditions on Model Fig. 3. None of these affect our qualitative conclusions.

The parameters that would have a more significant effect on the model relate to the term that defines the interfacial energy on chromosomes in Eq. (4): $A\left[1+\frac{b(w-w_0)^2}{w_0^2}\right]\varsigma e_{SS}$. Specifically, the interfacial energy parameter $\boldsymbol{\zeta}$, the thickness penalty parameter $b$, and the optimal thickness parameter $w_0$. As discussed below, changes in these parameters will lead to changes in the relationship between $\alpha$, $V_c$ and the number of synapsed chromosomes (Model Fig. 4). However, the qualitative conclusions of the model are all unchanged.

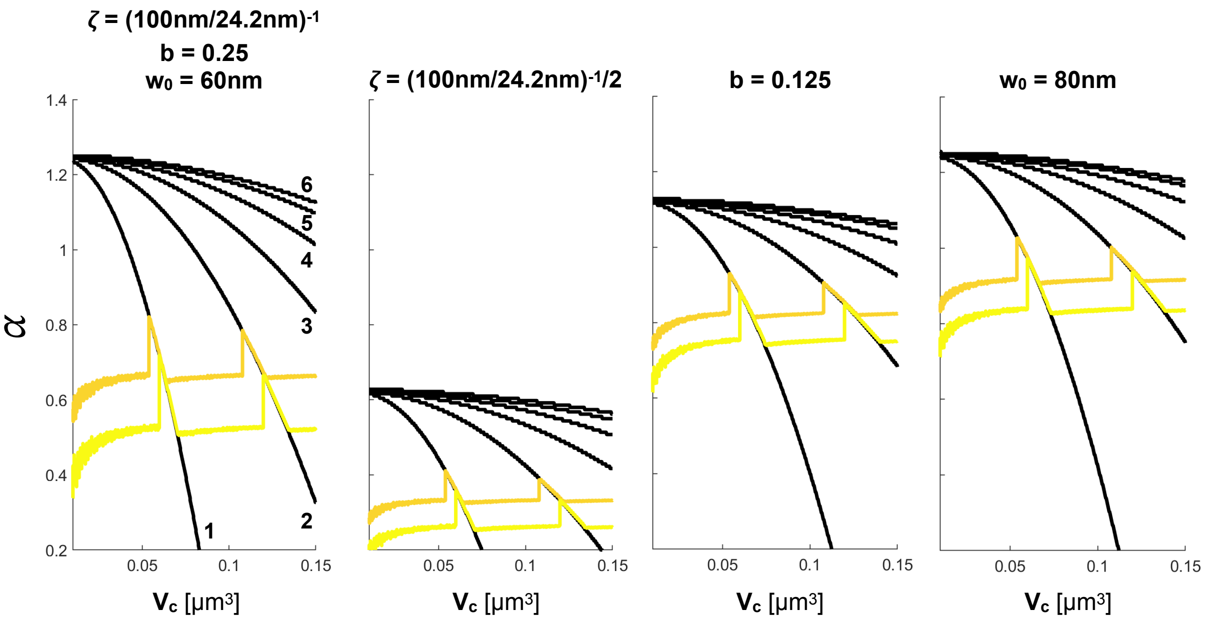


**Model Figure 4: Robustness analysis.** Left, contour plot using the same parameters as in Model Fig. 3 is shown for reference. The three plots to the right alter one of the parameters by 2-fold, as indicated above each plot.

##### 4.1 The interfacial energy parameter *ζ*

The parameter *ζ* estimates the loss of binding energy for an SC-CR molecule when it is exposed at the interface. We have estimated *ζ* based on the inter-axes distance (100nm) and the ladder spacing (24.2nm) to be 1/100nm/24.2nm. Halving this parameter would cause $\alpha$ to be halved (Model Fig. 4). However, that does not qualitatively change any of our findings. Moreover, since $\alpha$ is in fact arbitrary unit, it could be argued that it should be set to 1 in the wild-type setting, with mutant scenarios presented as relative to the wild-type value.

##### 4.2 The thickness penalty parameter b

Changing this penalty function can have a drastic effect on the contour plots (Model Fig. 4). The shift can affect both $\alpha$ and $V_c$. While we have no way to experimentally constrain this parameter, it does not affect any of the qualitative conclusions.

##### 4.3 The optimal thickness parameter $w_0$

Changing the optimal thickness $w_0$ can have a drastic effect on the contour plots (Model Fig. 4). As with the thickness penalty parameter, the shift can affect both $\alpha$ and $V_c$. These changes do not have any bearing on our qualitative conclusions.

While we cannot directly measure $w_0$, our measurements in mutant scenarios (such as *him-3*$^{KK170\text{-}171EE}$) allow to estimate an upper limit based on the appearance of multi-lamellar SC-CR. This consideration indicates at least a ~2-fold tolerance relative to the typical wild-type thickness (30-40nm; (Köhler et al. 2017, 2020)). Future work in experimental systems that allow over-expression of SC-CR components (Voelkel-Meiman et al. 2012) could further constrain this number.
